# A palmitoylation code controls PI4KIIIα complex formation and PI(4,5)P_2_ homeostasis at the plasma membrane

**DOI:** 10.1101/2021.06.14.447954

**Authors:** Alex G. Batrouni, Nirmalya Bag, Henry Phan, Barbara A. Baird, Jeremy M. Baskin

## Abstract

PI4KIIIα is the major enzyme responsible for generating the phosphoinositide PI(4)P at the plasma membrane. This lipid kinase forms two multicomponent complexes, both including a palmitoylated anchor, EFR3. Whereas both PI4KIIIα complexes support production of PI(4)P, the distinct functions of each complex and mechanisms underlying the interplay between them remain unknown. Here, we present roles for differential palmitoylation patterns within a tri-Cys motif in EFR3B (Cys5/Cys7/Cys8) in controlling the distribution of PI4KIIIα between these two complexes at the plasma membrane and corresponding functions in phosphoinositide homeostasis. Spacing of palmitoyl groups within three doubly palmitoylated EFR3B “lipoforms” affects both its interactions with TMEM150A, a transmembrane protein governing formation of a PI4KIIIα complex functioning in rapid PI(4,5)P_2_ resynthesis following PLC signaling, and its partitioning within liquid-ordered and -disordered regions of the plasma membrane. This work identifies a palmitoylation code in controlling protein–protein and protein–lipid interactions affecting a plasma membrane-resident lipid biosynthetic pathway.

**SUMMARY STATEMENT:** Different palmitoylation patterns on a lipid kinase adaptor protein control partitioning of the kinase between two spatiotemporally and functionally distinct complexes within the plasma membrane.

## INTRODUCTION

Phosphoinositides (PIPs) are a class of low abundance phospholipids that share the common biosynthetic precursor phosphatidylinositol (PI). Despite their scarcity, PIPs play crucial roles in most aspects of cellular life, from signal transduction to cell proliferation and recruitment of proteins to membranes in a spatially and temporally regulated manner^[1]^. Their defining structural feature is a *myo*-inositol headgroup that can be phosphorylated and dephosphorylated by lipid kinases and phosphatases at positions 3, 4, and 5 on the inositol ring, generating a total of seven distinct PIP species^[1]^. The most abundant member of this lipid family, phosphatidylinositol 4-phosphate (PI(4)P), is located in multiple organelle membranes and plays diverse roles in the cell. At the plasma membrane (PM), it serves as a biosynthetic precursor in the major pathway to produce PI(4,5)P_2_ and PI(3,4,5)P_3_. Beyond this biosynthetic role, PI(4)P at the PM has many functions^[2]^, including regulating non-vesicular transport and homeostasis of phosphatidylserine^[3]^ and other lipids^[4]^, modulating the activity of numerous ion channels^[5]^, serving as a substrate for certain GPCRs^[6,7]^, and, via its strongly anionic charge, helping to establish PM identity^[5]^.

PI(4)P is synthesized by phosphorylation of PI by PI 4-kinases. Of the four PI 4-kinase isoforms present in mammalian cells, a single enzyme, PI4KIIIα, encoded by the gene PI4KA, is responsible for the overwhelming majority of the PI(4)P pool at the PM, with other non-redundant PI4K isoforms responsible for producing biochemically distinct PI(4)P pools on intracellular membranes. Speaking to the vital importance of the PM pool of PI(4)P, PI4KIIIα is essential for life in all model organisms tested, including mice, fruit flies, and yeast^[8–10]^. PI4KIIIα function is important to human health, as mutations to some of its non-catalytic accessory proteins, described below, cause a variety of disorders, ranging from hypomyelination^[11–13]^ to severe immunodeficiencies^[14]^ and congenital intestinal obstructions^[12,15]^. Further, several types of viruses hijack this enzyme during infection to remodel internal cellular membranes in support of viral replication^[16,17]^. Given these numerous and important (patho)physiological functions of the PM pool of PI(4)P, the regulation of its levels is critically important.

Because PI4KIIIα is a soluble protein lacking membrane-anchoring motifs^[8]^, its recruitment to the PM to access its lipid substrate, PI, requires the action of several additional protein factors. In 2001, the six-pass transmembrane protein suppressor of four-kinase 1 (Sfk1) was first identified in yeast as a factor whose overexpression partially suppressed a temperature-sensitive mutant of Stt4, the PI4KIIIα ortholog in that organism^[18]^. Subsequently, two essential proteins, Efr3 and Ypp1, were characterized to form a complex with Stt4 that recruits it to the PM^[19]^, though the relationship of this complex to Sfk1 remains unclear. The mammalian orthologs of Efr3 and Ypp1, EFR3A/EFR3B and TTC7A/TTC7B, respectively, perform similar roles in recruiting PI4KIIIα to the PM^[8]^, with the mammalian PI4KIIIα complex containing an additional factor, FAM126A (also known as hyccin)/FAM126B^[11,13]^.

A combination of structural, biochemical, and imaging studies has led to a working model for the EFR3–TTC7–FAM126–PI4KIIIα complex, which we will term Complex I (**Fig. 1A, left**) ^[13,20–22]^. EFR3 constitutively localizes to the PM by electrostatic interactions from a basic patch on its folded α-helical N-terminal domain, and, critically, palmitoylation of an N-terminal Cys-rich motif^[8]^. The unstructured C-terminal tail of EFR3 binds to TTC7^[23]^, which is part of a globular hexameric complex comprising two copies each of TTC7, FAM126, and PI4KIIIα^[21,22]^. As for a potential role of Sfk1 in mammals, the 271-residue, six-pass transmembrane protein TMEM150A was identified as its functional ortholog in 2015^[20]^. This study reported that PI4KIIIα can exist in a second complex, comprising EFR3, TMEM150A, and PI4KIIIα, which we will term Complex II (**Fig. 1A, right**) and which, when overexpressed, accelerated PI(4,5)P_2_ resynthesis after acute depletion, presumably via production of PI(4)P, the substrate for type I PIP5K enzymes that synthesize PI(4,5)P_2_. Interestingly, the TMEM150A-containing Complex II was only observed to form downstream of the TTC7- and FAM126-containing Complex I. This model suggests a degree of interplay between these two complexes, wherein PI4KIIIα and EFR3 are handed off from TTC7/FAM126 to TMEM150A.

**Figure 1.**
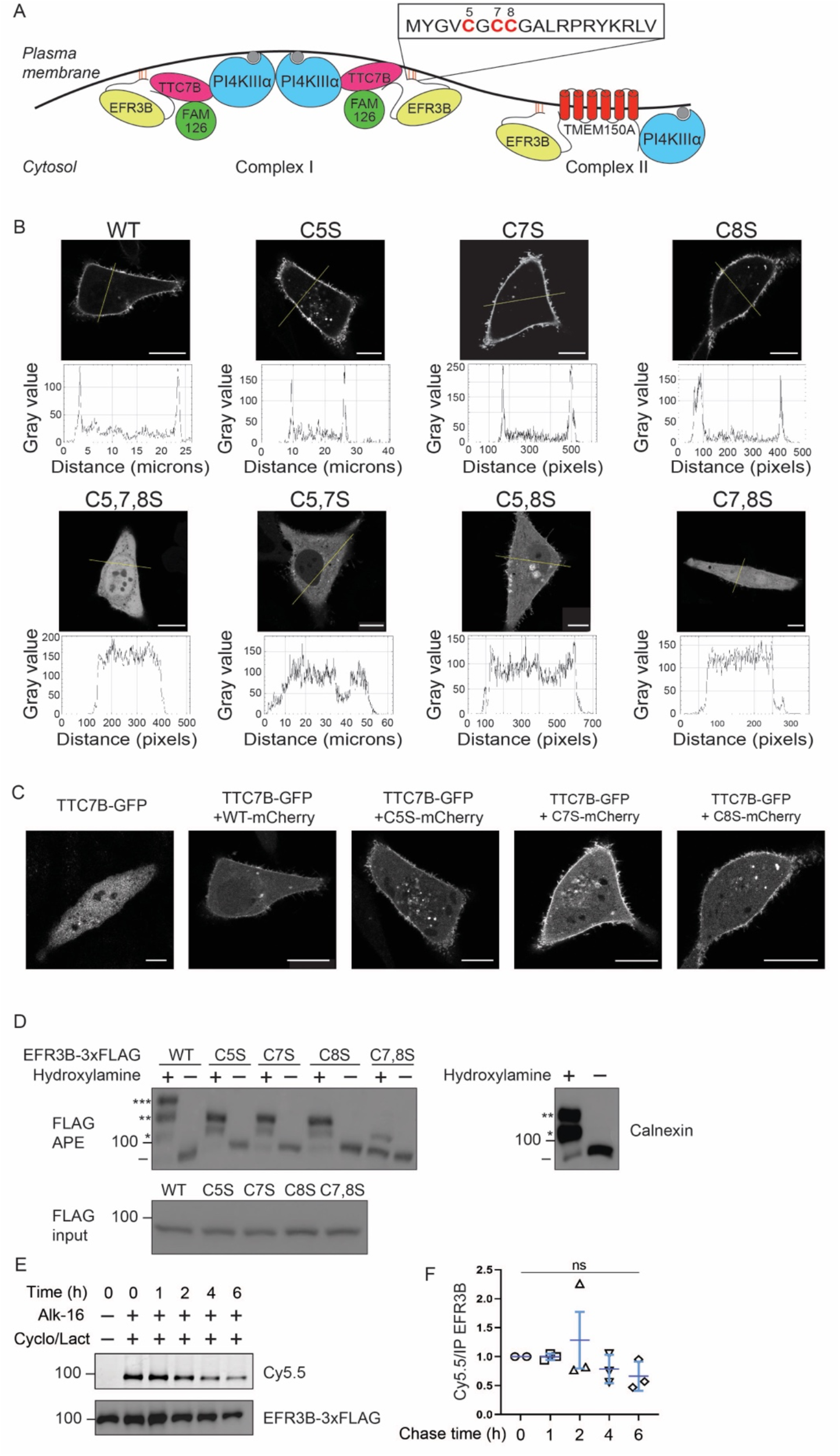
Dual palmitoylation of EFR3B is required for its PM localization. (A) Diagram illustrating the current understanding of the assembly of the PI4KIIIα into two complexes (Complex I, left, and Complex II, right) at the PM. The N-terminal sequence of EFR3B, including the Cys residues within the palmitoylation motif, is highlighted. (B) Confocal microscopy images of EFR3B-mCherry mutants expressed in HeLa cells: WT, single Cys mutants (CxS: C5S, C7S, and C8S), double Cys (CxxS: C5,7S, C5,8S, and C7,8S), and the triple Cys mutant (C5,7,8S). Traces underneath each image indicate fluorescence intensity along the line indicated in the image, to illustrate the presence/absence of discernable PM fluorescence. Scale bars, 15 μm. (C) Demonstration that all EFR3B-mCherry single Cys mutants are capable of recruiting TTC7B-GFP to the PM similarly to WT EFR3B-mCherry. Shown is TTC7B-GFP fluorescence from cells co-transfected with TTC7B-GFP and the indicated EFR3B-mCherry construct. Scale bars, 15 μm. (D) APE assay to quantify the extent of palmitoylation of various EFR3B-3xFLAG mutants, with calnexin used a positive control. Lysates containing EFR3B-3xFLAG were subjected to acyl-PEG exchange, with 5 kD PEG replacing each palmitoyl group. The FLAG and calnexin APE blot shows the result of the APE assay, with EFR3B or calnexin species with zero, one, two, or three PEG groups indicated by the presence of −, *, **, or *** respectively. A sample of lysate prior to APE was analyzed to show input (FLAG, input). (E–F) Bioorthogonal metabolic labeling pulse-chase experiment with alk-16 showing that EFR3B-3xFLAG palmitoylation is stable over several hours. Cells metabolically labeled overnight with alk-16 were chased in regular media for 0–6 h in the presence of cycloheximide and lactacystin to prevent protein synthesis and degradation (Cyclo/Lact), followed by lysis, FLAG immunoprecipitation, click chemistry tagging with Cy5.5-azide, and SDS-PAGE analysis. Shown in (E) are in-gel fluorescence (Cy5.5, to indicate extent of palmitoylation) and Western blot (to indicate total EFR3B-3xFLAG). Quantification of the data are shown in (F), with Cy5.5 fluorescence normalized to EFR3B-3xFLAG intensity in the Western blot (n=3). Statistical significance was determined using Student’s t-test. ns, not significant.

It remains unclear why cells contain two distinct PI4KIIIα-containing complexes within the same membrane to generate the same lipid product, PI(4)P. Critically, the mechanisms by which cells control the balance between both complexes have not been investigated. Given their compositional differences, it is conceivable that Complexes I and II are stabilized in the PM by different modes of interactions. Though Complex I is primarily anchored to the PM by EFR3, TTC7 and PI4KIIIα engage in secondary, stabilizing electrostatic interactions with anionic lipids in the inner leaflet of the PM via a handful of conserved, positively charged residues on their surfaces^[21]^. By contrast, Complex II does not contain TTC7 but includes TMEM150A, and presumably both EFR3 and TMEM150A play important roles in anchoring PI4KIIIα to the PM. The differential mechanisms by which PI4KIIIα is anchored in the PM could form the basis for differences in assembly and functional properties between the two complexes.

The palmitoylated protein EFR3 is the only non-catalytic protein shared between Complexes I and II. We reasoned that a detailed understanding of the how it associates with membranes might reveal mechanisms underlying the interplay between the two PI4KIIIα complexes. Palmitoylation, or S-acylation, is the post-translational modification of Cys residues as fatty acyl thioesters, typically with 16-carbon, saturated palmitoyl groups. Palmitoylation of target proteins induces their anchoring within membranes, and its reversible nature makes this a powerful mechanism to regulate protein localization, degradation, and function^[24–26]^. Palmitoylation of Cys-rich motifs within proteins can mediate partitioning of such proteins to liquid-ordered like (Lo-like) membrane domains, which are enriched in lipids with saturated tails^[27]^. Interestingly, PI(4,5)P_2_ synthesis was recently proposed to occur in these domains^[28]^, though the mechanisms of Lo or the complementary liquid-disordered (Ld)-like membrane domain compartmentalization of PI(4,5)P_2_ synthesis are not fully understood.

The N-terminus of EFR3B, one of two mammalian EFR3 proteins implicated in PI4KIIIα function, contains three Cys residues that can be palmitoylated (C5, C7 and C8)^[8,20]^. Mutation of all three to Ser abolished the PM localization of EFR3B, but the contributions of palmitoylation at individual Cys residues within this motif to EFR3 localization and PI4KIIIα function have not been examined^[8]^. Due to the reversible nature of palmitoylation and the potential for changes to the extent and pattern of palmitoylation on EFR3 to impact its interactions within the two PI4KIIIα complexes, we hypothesized that palmitoylation of EFR3B could represent a regulatory mechanism governing PI4KIIIα recruitment to specific membrane subregions within distinct complexes.

In this study, we have undertaken a detailed examination of the role of EFR3 palmitoylation as a mechanism regulating the assembly of PI4KIIIα into two spatiotemporally and functionally distinct complexes at the PM. A key aspect of our study was understanding how interactions of EFR3 with TMEM150A modulate the biophysical and catalytic properties of the PI4KIIIα complex in the PM. Using a combination of biochemistry, chemical biology, and biophysical imaging techniques, we found that the overwhelming majority of EFR3B molecules are modified with two or three palmitoyl groups, and that any two of the three Cys residues in EFR3B are sufficient for localization to the PM. Further, we found that EFR3B palmitoylation is stable over the course of multiple hours, giving rise to several “lipoforms” of EFR3B, i.e., distinct pools of EFR3B with different palmitoylation patterns, that coexist in the PM. Intriguingly, we found that TMEM150A preferentially interacts with the EFR3B lipoform palmitoylated on the adjacent Cys residues at positions 7 and 8 (EFR3B(7,8-palm)). Using an in situ, imaging-based detergent resistant membrane (iDRM) assay, we revealed that Complex II preferentially localizes to Ld-like regions of the PM, suggesting a Ld-like specific function for PI(4,5)P_2_ synthesis. Finally, quantification of PI(4,5)P_2_ resynthesis following acute depletion under conditions favoring the formation of Complex I or II solidifies a role for Complex II as most efficiently producing PI(4)P for this functional process.

Based on these data, we propose a comprehensive framework explaining PI4KIIIα association with and function in the PM. First, PI4KIIIα is recruited to the PM by EFR3B^[8,13,20]^ within Complex I, which localizes to Lo-like regions of the PM. Subsequently, the preferential interaction of TMEM150A with EFR3B(7,8-palm) associated with PI4KIIIα leads to formation of Complex II, which localizes to Ld-like regions of the PM. The pool of PI4KIIIα associating with other EFR3B largely remains in Complex I. This model provides a basis for understanding how cells maintain two spatiotemporally and functionally distinct PI4KIIIα complexes to facilitate synthesis of PI(4)P within different subdomains of the PM.

## RESULTS

### Two of the three Cys residues of EFR3B are required for its PM localization

EFR3B contains three Cys residues (C5, C7, and C8) within an N-terminal palmitoylation motif (**Fig. 1A**). It was previously shown that mutation of all three N-terminal Cys residues of EFR3B (and all four within its homolog, EFR3A), completely abolishes its PM localization[8]. Yet, the extent of palmitoylation of wild-type EFR3B, and the palmitoylation requirements for PM localization and function in the PI4KIIIα complex, remained unknown. Therefore, we systematically evaluated the link between the palmitoylation status and PM localization of EFR3B. We generated all permutations of single and double Cys to Ser mutants in mCherry-tagged EFR3B. Using confocal microscopy imaging, we found that mutation of any single Cys residue (i.e., C5S, C7S and C8S mutants, collectively referred to as CxS or “dual-palm” mutants) had no impact on EFR3B-mCherry anchoring to the PM (**Fig. 1B, top panel**). However, mutation of any pair of Cys residues (i.e., “single-palm” mutants) resulted in the mislocalization of the overwhelming bulk of EFR3B-mCherry to the cytosol (**Fig. 1B, bottom panel**). The triple Cys mutant (C5,7,8S) was completely absent from the PM as previously reported (**Fig. 1B**)^[8]^. Therefore, two of the three Cys residues within the EFR3B N-terminal motif are minimally required for its anchoring to the PM.

We next tested the ability of the CxS/dual-palm mutants to recruit TTC7B to the PM as a proxy for formation of Complex I. It was previously reported that overexpression of WT EFR3B is necessary to recruit transiently transfected TTC7B to the PM^[8]^. Here, we examined the localization of these proteins by confocal microscopy in cells co-expressing TTC7B-GFP and either WT or a Cys mutant form of EFR3B-mCherry. We found that all three CxS mutants were also able to recruit TTC7B as efficiently as WT EFR3B (**Fig. 1C**).

The above results indicate that any two of the three N-terminal Cys residues of EFR3B are sufficient for its localization and capacity to recruit TTC7B at the PM. This led us to ask whether all three N-terminal Cys residues of WT EFR3B are actually palmitoylated. We used the acyl PEG exchange (APE) assay^[29]^ to assess the degree of palmitoylation in WT and CxS mutants of EFR3B. APE allows for tagging of palmitoylated Cys residues with a mass-tag (5 kDa PEG in this case), resulting in a molecular weight shift that can be visualized by SDS-PAGE/Western blot. We found that WT EFR3B was roughly split equally between double and triple palmitoylated forms, with very minimal amounts of single or non-palmitoylated protein observed (**Fig. 1D**). All three EFR3B CxS mutants were nearly completely doubly palmitoylated. By contrast, a representative CxxS (“single-palm”) mutant, C7,8S, was examined and found to exist predominantly in its non-palmitoylated form, with a minority singly palmitoylated form. Combined with the imaging results, these data confirm both that dual palmitoylation of EFR3B is required for its PM localization and that two Cys residues within this motif result in its efficient, i.e., near-stoichiometric, palmitoylation. Furthermore, we found that co-expression of TMEM150A with EFR3B had no effect on the palmitoylation status of EFR3B (**Fig. S1**).

### EFR3B palmitoylation is stable

Palmitoylation is generally known to be reversible post-translational modification that can, in certain contexts, dynamically regulate protein-membrane and protein-protein interactions^[27]^. We next investigated the dynamics of EFR3B palmitoylation, using metabolic labeling of cells expressing FLAG-tagged EFR3B with a bioorthogonal, alkyne-containing palmitate analog (alk-16)^[30]^. To evaluate the stability of palmitoylation, cells were then chased with regular media in the presence of cycloheximide and lactacystin (inhibitors of protein synthesis and the proteasome, respectively) for up to six hours, followed by lysis, FLAG immunoprecipitation (IP), click chemistry tagging to label alk-16-containing proteins with a Cy5.5-azide probe, and SDS-PAGE analysis and either in-gel fluorescence to assess palmitoylation, or Western blot for FLAG to detect total EFR3B-FLAG protein. In contrast to other proteins that display very rapid palmitoylation/depalmitoylation dynamics^[31]^, we found that palmitoylation of EFR3B is stable over the course of several hours (**Fig 1E and 1F**). Collectively, these studies of EFR3B palmitoylation indicate that the vast majority of WT EFR3B is doubly or triply palmitoylated and that this lipidation, responsible for its PM localization, is stable on the multi-hour timescale.

### Interactions between EFR3B and TMEM150A modulate their respective membrane dynamics

We next explored how palmitoylation of EFR3B impacts its interactions with TMEM150A, the only other constitutively PM-localized component involved in PI4KIIIα recruitment^[20]^. Previously, EFR3B and TMEM150A were shown to interact by co-immunoprecipitation (co-IP)^[20]^, though no discrete binding sites on either protein were identified. Therefore, the mode of interaction may not rely entirely on protein-protein binding. The interactions of PM-localized proteins are often regulated by the collective biophysical properties of the PM organization itself. For example, lipid-dependent, dynamic partitioning into liquid ordered-like (Lo-like) nanodomains helps to concentrate selected proteins to facilitate their interactions^[32]^. In this context, it is worth noting that palmitoylation is one of the primary mechanisms by which PM proteins can partition into Lo-like nanodomains^[27]^. We hypothesized that EFR3B and TMEM150A could participate in protein-protein and protein-lipid interactions at the PM, mediated by dynamic organization of this membrane, and that the palmitoylation status of EFR3B could play a role in promoting these interactions. To test this hypothesis, we characterized the PM dynamics and organization of EFR3B and TMEM150A both alone and together.

We performed fluorescence recovery after photobleaching (FRAP) experiments on the ventral PM of a rat basophilic leukemia cell line (RBL-2H3) expressing a fluorescently tagged protein to quantitively assess the protein’s diffusion rate and the mobile fraction. We found that the average recovery time (*t*_1/2_) of WT EFR3B-GFP was 16±5 s (**Fig. 2A**), which was similar to other PM inner leaflet probes in RBL-2H3 cells measured under identical instrumental conditions^[33]^. Likewise, the mobile fraction was >80%, indicating that the diffusing pool represents the majority of EFR3B molecules. Because WT EFR3B exists as a mixture of triply and doubly palmitoylated species (**Fig. 1D**), we also examined if the observed diffusion properties are dictated by one of these species. We performed FRAP experiments on all three fluorescently tagged CxS (dual-palm) mutants and found that the recovery time and mobile fraction of the dual-palm mutants were the same as WT EFR3B (**Figs. 2E** and **S2**). We note that, among these mutant constructs, GFP-tagged C5S and C8S localized correctly (i.e., identically to WT), but GFP-tagged C7S exhibited unusual puncta at the PM. Because its mCherry and mNeonGreen (mNG) fusion protein counterparts did not show such features, we concluded that the punctate structures formed by the GFP-tagged version were artifactual, therefore we performed FRAP measurements on mNG-tagged C7S. Overall, these data confirm that the diffusion properties of triply and doubly palmitoylated species are the same.

**Figure 2.**
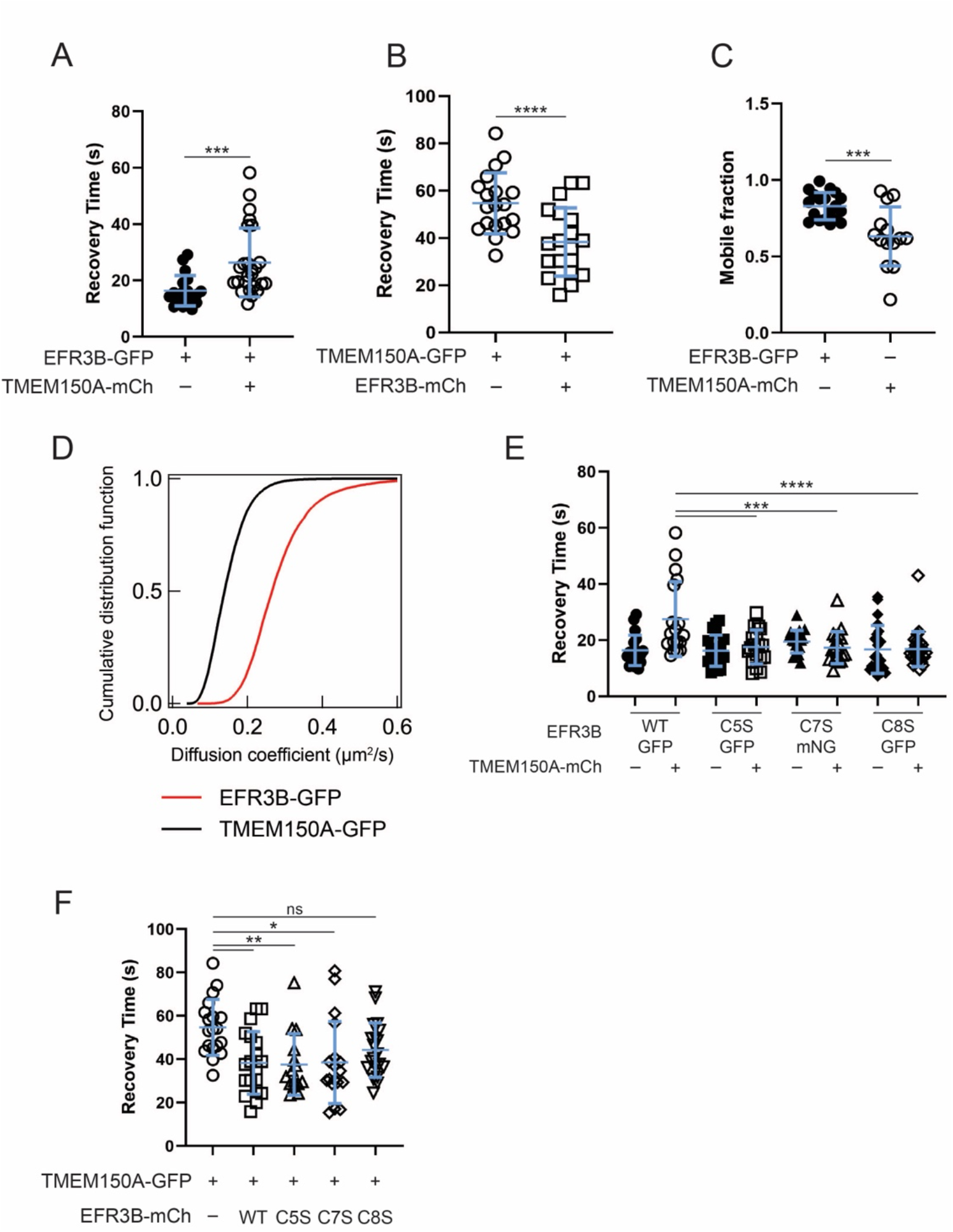
FRAP studies reveal that EFR3B interacts biophysically with TMEM150A and that the membrane dynamics of WT and CxS EFR3B mutants are similar. (A) FRAP recovery times (*t_1/2_*) of EFR3B-GFP in the absence and presence of TMEM150A-mCherry. The increase in the recovery time of EFR3B in presence of TMEM150A suggests an interaction between the two proteins. (B) FRAP recovery times (*t_1/2_*) of TMEM150A-GFP in absence and presence of EFR3B-mCherry. Note that the recovery time of TMEM150A is reduced in presence of EFR3B to match that of EFR3B in (A). (C) Mobile fraction of EFR3B-GFP or TMEM150A-GFP when expressed alone as evaluated by FRAP. (D) Diffusion coefficients of EFR3B-GFP and TMEM150A-GFP measured by ImFCS. (E) Recovery times (*t_1/2_*) of WT EFR3B-GFP and individual EFR3B-GFP/mNG CxS mutants in the absence and presence of TMEM150A-mCherry. Note that the CxS mutations have no effect on the membrane diffusion of EFR3B and are insensitive to the presence of TMEM150A. (F) Recovery time (*t_1/2_*) of TMEM150A-GFP expressed alone or in presence of WT EFR3B-mCherry or each of the three EFR3B-mCherry CxS mutants. Note that the CxS mutants have the same effect on TMEM150A-GFP recovery time as WT EFR3B. Statistical significance was determined using a one-way ANOVA with Tukey-Kramer post-hoc test. n=14–27; *, p<0.05; **, p<0.01; ***, p<0.005; ****, p<0.001, ns = not significant.

In contrast to EFR3B-GFP, the transmembrane protein TMEM150A-GFP showed a much slower recovery (*t*_1/2_ = 55±13 s), with a smaller mobile fraction (~60%) (**Figs. 2B–C**), consistent with studies indicating that the diffusion of transmembrane proteins is generally slower than lipid-anchored proteins^[34,35]^. As a control experiment, we found that the FRAP recovery properties of TMEM150A-GFP were similar to its homolog TMEM150B-GFP, which is structurally very similar but not involved in formation of the PI4KIIIα complex^[20]^ (**Fig. S3**). However, the diffusion of the TMEM150 proteins is much slower than other multipass transmembrane proteins measured previously^[33]^. For example, the *t*_1/2_ of the seven-pass immunoglobulin E (IgE) receptor, FcεRI, was 18±4 s when measured on an identical instrumental setup^[33]^, indicating faster diffusion than TMEM150A-GFP. Such slow diffusion of TMEM150A-GFP is not due to clustering, because coexpression of TMEM150A-GFP and TMEM150A-mCherry did not show further reductions in diffusion rates or mobile fraction (**Fig. S4**), indicating that the observed slow diffusion is likely not due to overexpression artifacts. Therefore, TMEM150A-GFP may be associated with other yet unknown membrane components that significantly slow down its diffusion. We also employed Imaging Fluorescence Correlation Spectroscopy (ImFCS) on RBL-2H3 cells expressing EFR3B-GFP or TMEM150A-GFP to measure their diffusion coefficients (*D*) with very high precision^[33,36,37]^. The cumulative distributions constructed from ~10,000 *D* values for each of the probes are compared in **Fig. 2D**. These experiments confirm the slower diffusion of TMEM150A-GFP relative to EFR3B-GFP, with average *D* values of 0.15±0.05 μm^2^/s and 0.28±0.09 μm^2^/s respectively.

Interestingly, the FRAP characteristics of both proteins change in the presence of the other. We observed an increase in the *t*_1/2_ (27±13 s) of EFR3B-GFP in the presence of TMEM150A-mCherry, with unaltered mobile fraction (**Fig. 2A**). This result suggests that the mobile population of EFR3B interacts with TMEM150A leading to a slowing of diffusion, consistent with the co-IP results reported previously^[20]^. We observed consistent changes in the diffusion behavior of TMEM150A-GFP co-expressed with EFR3B-mCherry (**Fig 2B**). The *t*_1/2_ of TMEM150A-GFP in the presence of EFR3B-mCherry decreased significantly (*t*_1/2_ = 38±14 s), and the mobile fraction was again unaltered. This result indicates that associating with EFR3B-mCherry is accompanied by liberating TMEM150A-GFP from its associations with other membrane components as speculated above. Notably, the *t*_1/2_ values for EFR3B-GFP and TMEM150A-GFP when co-overexpressed were close to one another (i.e., 27±13 s and 38±14 s respectively). Because the respective mobile fractions of the two proteins did not change in the presence of the other but the recovery time did, it is likely that the majority of the mobile populations of these proteins interact with each other.

To test if the diffusion changes of fluorescently tagged EFR3B and TMEM150A in the presence of each other are specific, we conducted a series of additional control experiments. First, we measured the diffusion of TMEM150A-GFP co-expressed with Lyn-mRFP, a PM-localized inner leaflet, lipid-anchored kinase. Here, the *t*_1/2_ and mobile fraction of TMEM150A-GFP remained unaltered, indicating that there are no non-specific interactions between Lyn and TMEM150A (**Fig. S5**). This scenario is similar to the unaltered membrane dynamics of TMEM150A-GFP in the presence of TMEM150A-mCherry, further suggesting that the faster diffusion of TMEM150A in the presence of EFR3B is the result of specific interactions. We performed reciprocal experiments to test if EFR3B impacts the diffusion of other transmembrane proteins. For this, we measured diffusion of a YFP-tagged muscarinic M1 receptor (M1R), a seven-pass GPCR, with or without co-overexpressed EFR3B-mCherry. The *t*_1/2_ of M1R-YFP was ~23±5 s in both conditions along with unaltered mobile fraction (**Fig. S6**). These observations establish the specificity of the EFR3B–TMEM150A interaction as evaluated by FRAP.

To assess the importance of EFR3B palmitoylation in the EFR3B–TMEM150A interaction, we monitored the diffusion changes of TMEM150A-GFP in the presence of the dual-palm EFR3B mutants. Co-overexpression of mCherry-tagged CxS EFR3B species had a largely similar effect to that of WT EFR3B-mCherry, also decreasing the *t*_1/2_ of TMEM150A-GFP, indicating that the doubly palmitoylated species are capable of interacting with TMEM150A-GFP (**Fig. 2F**). By contrast, the diffusion of GFP- or mNG-tagged CxS species, unlike WT EFR3B, did not show reduced diffusion (i.e., *t*_1/2_ was unchanged) when co-overexpressed with TMEM150A-mCherry (**Fig. 2E**).

As described earlier, we note that mNG-tagged EFR3B(C7S) was used because of clustering observed with the GFP-tagged version of this construct, but not the mNG- or mCherry-tagged versions. FRAP experiments with the various EFR3B(C7S) constructs indicated that the GFP fusion displayed much slower diffusion kinetics than the other two GFP-tagged CxS constructs, whereas the mNG- and mCherry-tagged C7S construct exhibited similar diffusion to the GFP-tagged CxS constructs (**Fig. S7**). Collectively, these results suggest that the dual-palm mutants of EFR3B can dissociate TMEM150A from its low-diffusion assembly with other membrane components, but that these mutants likely do not remain associated with TMEM150A for a sufficiently long period of time for this interaction to have an impact on their own mean diffusion speed.

### TMEM150A preferentially interacts with certain EFR3B lipoforms within liquid-disordered (Ld)-like membrane regions

Palmitoylation is one of the primary mechanisms by which PM-localized proteins preferentially partition in liquid ordered like (Lo-like) nanodomains^[38]^. These nanodomains function to provide optimal spatial compartmentalization for productive interactions between proteins. We recently developed an in situ, imaging-based detergent resistant membrane (iDRM) assay (**Fig. 3A**) to quantify relative preference of a protein to Lo-like nanodomains^[33]^. The principle behind this assay is that Lo-preferring, fluorescently tagged proteins within the PM are more resistant to mild treatment with the detergent Triton X-100 (TX-100) than proteins in Ld-like nanodomains. Therefore, a higher fraction of such Lo-preferring proteins will remain in the PM of intact cells upon TX-100 treatment (**Fig. 3A**). The key metric obtained from iDRM experiments is the resistance (*R*) value, defined as the ratio of fluorescence in TX-100-treated cells to that in vehicle-treated cells. A higher *R* value represents a stronger interaction between the target protein and Lo-like nanodomain, resulting in more retention of the protein in the Lo-like nanodomain. A lower *R* value suggests a weaker interaction, as more protein is extracted from the PM by the detergent wash. Note that other factors may lead to detergent resistance of proteolipid and transmembrane probes regardless of domain partitioning, such as interactions with the actin cytoskeleton and other detergent-resistant proteins.

**Figure 3.**
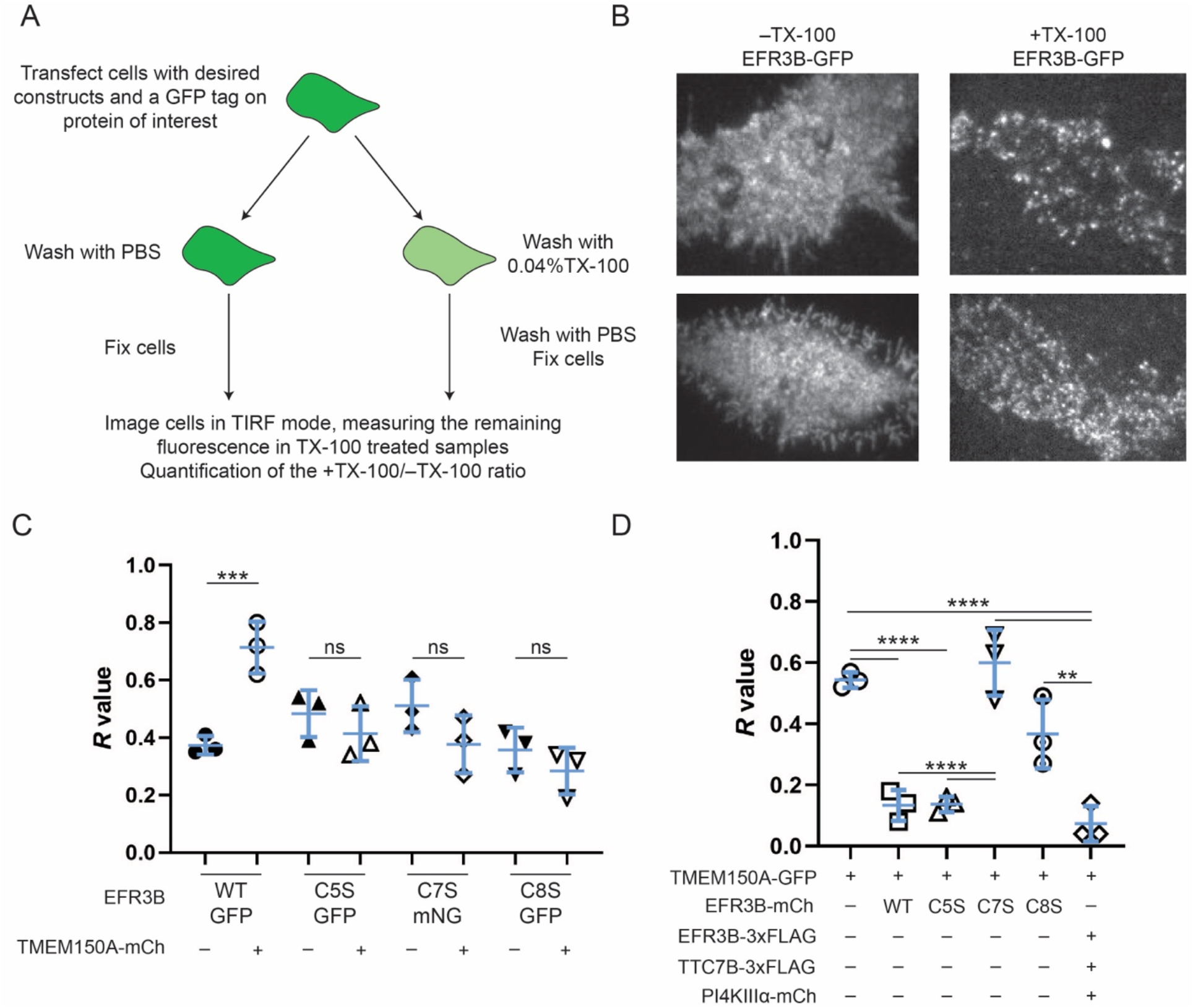
TMEM150A preferentially interacts with EFR3B(C5S) to form the PI4KIIIα complex. (A) Diagram illustrating the workflow of the imaging detergent resistant membrane (iDRM) assay. All DRM experiments were performed in RBL-2H3 cells. (B) Representative images of RBL-2H3 cells expressing EFR3B-GFP treated with PBS or 0.04% TX-100 in the iDRM assay. (C) Quantification of iDRM experiments on EFR3B-GFP (WT or the indicated CxS mutants) in the presence and absence of TMEM150A-mCherry, with the *R* value (fraction of retained fluorescence) plotted. Note that WT EFR3B exhibits a strong increase in detergent resistance in presence of TMEM150A-mCherry. However, the EFR3B(CxS) palmitoylation mutants do not. (D) Quantification of iDRM experiments on TMEM150A-GFP in presence of EFR3B-mCherry (WT or indicated CxS mutants), or the PI4KIIIα complex containing EFR3B-3xFLAG, TTC7B-3xFLAG, PI4KIIIα-mCherry. For the iDRM experiments, each data point represents the mean between 30 +TX-100 and 30 –TX-100 cells. Statistical significance was determined using a one-way ANOVA with Tukey-Kramer post-hoc test. n=3; **, p<0.01; ***, p<0.005; ****, p<0.001.

We performed iDRM experiments using the same (co)-overexpression conditions as the previous FRAP experiments. Representative images of cells expressing EFR3B-GFP washed with PBS or 0.04% TX-100 (**Fig. 3B**) show a modest amount of detergent resistance, yielding an *R* value of 0.37±0.03 (mean±SD) for this probe. The detergent resistance of EFR3B-GFP strongly increased (*R* = 0.71±0.07) when co-expressed with TMEM150A-mCherry. Interestingly, this was not the case for any of the dual-palm mutants (**Fig 3C**). These observations are analogous to the increased *t*_1/2_ of WT EFR3B, but not for the EFR3B-CxS mutants, in the presence of TMEM150A as measured by FRAP (**Fig. 2E**). The changes in *R* (iDRM) and *t*_1/2_ (FRAP) values of WT EFR3B may be governed by an elevated Lo-partitioning of triple palmitoylated lipoform of EFR3B in this co-expression condition.

By monitoring TMEM150A-derived fluorescence instead of EFR3B fluorescence in iDRM studies, we found that TMEM150A-GFP was more detergent-resistant (*R* = 0.54±0.02) than EFR3B-GFP (**Fig. 3C and 3D**). Because we found that TMEM150A did not form homo-oligomers (**Fig. S3**), we reasoned that its detergent resistance may stem from its Lo-preference and/or heterotypic interactions with other proteins. Interestingly, TMEM150A-GFP became highly susceptible to detergent when co-expressed with EFR3B-mCherry (**Fig. 3D**, *R* = 0.13±0.04). This effect was replicated by co-expression with EFR3B(C5S)-mCherry (**Fig. 3D**, *R*= 0.14±0.02). Yet, strikingly, EFR3B(C7S)-mCherry mutant had no effect on the detergent resistance of TMEM150A-GFP (**Fig. 3D**, *R* = 0.60±0.09). EFR3B(C8S)-mCherry showed an intermediate effect on the detergent resistance of TMEM150A-GFP (*R* = 0.37±0.09; **Fig. 3D**).

Finally, we tested the detergent resistance of key components of Complexes I and II (**Fig. 1A**). The detergent resistance of GFP-PI4KIIIα was similar when cells were co-transfected with plasmids encoding the necessary components to assemble Complex I (EFR3B-FLAG, TTC7B-FLAG, and GFP-PI4KIIIα, *R* = 0.51±0.11) or Complex II (EFR3B-FLAG, TTC7B-FLAG, GFP-PI4KIIIα, and TMEM150A-mCherry, *R*= 0.55±0.07) (**Fig. S8**). Recent cryo-EM and HDX-MS structures of Complex I revealed extensive protein-protein interactions via the formation of a hexamer containing two copies each of TTC7B, FAM126A and PI4KIIIα (and presumably EFR3B)^[21,22]^. These strong protein-based interactions could explain the detergent resistance of GFP-PI4KIIIα as part of Complex I. Interestingly, when TMEM150A, the unique component of Complex II, was co-expressed with PI4KIIIα, EFR3B, and TTC7B, we found that it showed a very low extent of detergent resistance (*R* = 0.07±0.05) (**Fig. S8**), similar to its *R* value when TMEM150A-GFP was co-expressed with the WT or C5S versions of EFR3B (*R* = 0.13±0.04 and *R* = 0.14±0.02 respectively) (**Fig. 3D**). These results indicate that Complex II is likely more Ld-preferring or has fewer intermolecular interactions than Complex I. Additionally, because the *R* value of GFP-PI4KIIIα remained unaltered under co-overexpression conditions to form Complexes I and II, these results raise the possibility that both complexes may coexist in the PM and that PI4KIIIα occupancy of Complex I may predominate.

Our iDRM results suggest that the location of N-terminal palmitoylation of EFR3B is key to the formation of Complex II. In particular, they indicate that TMEM150A interacted more strongly with EFR3B palmitoylated at positions C7 and C8 (as occurs in the EFR3B(C5S) mutant). By contrast, we detected minimal interaction between TMEM150A and the EFR3B(C7S) mutant by iDRM, where the palmitoyl groups at C5 and C8 are separated by two residues. The interactions of TMEM150A and the EFR3B(C8S) mutant, which can be palmitoylated at positions C5 and C7 (separated by only one residue), showed an intermediate phenotype in the iDRM assay. Overall, the spacing between palmitoylation sites on EFR3B, as well as the extent of their modification, could modulate the partitioning of EFR3B, leading to preferential interaction of TMEM150A with both the triply palmitoylated and the C7/C8 doubly palmitoylated lipoforms of EFR3B.

### Interactions between EFR3B lipoforms and TMEM150A differentially impact the kinetics of PI(4,5)P_2_ synthesis

Finally, we investigated the functional significance of the preferential interaction between TMEM150A and the differentially palmitoylated forms of EFR3B. A key role for PI4KIIIα is to provide PI(4)P for subsequent phosphorylation by PIP5K enzymes to produce PI(4,5)P_2_, and demand for PI(4,5)P_2_ synthesis is particularly high following its transient depletion by Gq–PLC signaling. TMEM150A has previously been implicated in PI(4,5)P_2_ homeostasis^[20]^, and we sought to investigate the role of the EFR3B–TMEM150A interaction in this process.

We used an established system for acutely depleting PI(4,5)P_2_ and then permitting its resynthesis in the PM of HeLa cells using sequential treatment with a GPCR agonist and its antagonist. This system takes advantage of the activation of the muscarinic M1 receptor (M1R) by its agonist oxotremorine-M (oxo-M), which leads to downstream activation of PLCβ and hydrolysis of a major portion of PI(4,5)P_2_ in the PM to diacylglycerol (DAG) and inositol trisphosphate (IP_3_). This signaling event is reversed by addition of the M1R antagonist atropine, allowing resynthesis of PI(4,5)P_2_ from PI, via the action of PI4KIIIα and PIP5Ks. The reversible nature of this system allows for monitoring of the recovery kinetics of PI(4,5)P_2_ by co-expression of a PI(4,5)P_2_-binding fluorescent biosensor, iRFP-PH(PLCδ), which localizes to the PM when it is replete with PI(4,5)P_2_ and to the cytosol upon PI(4,5)P_2_ depletion (**Fig. 4A**).

**Figure 4.**
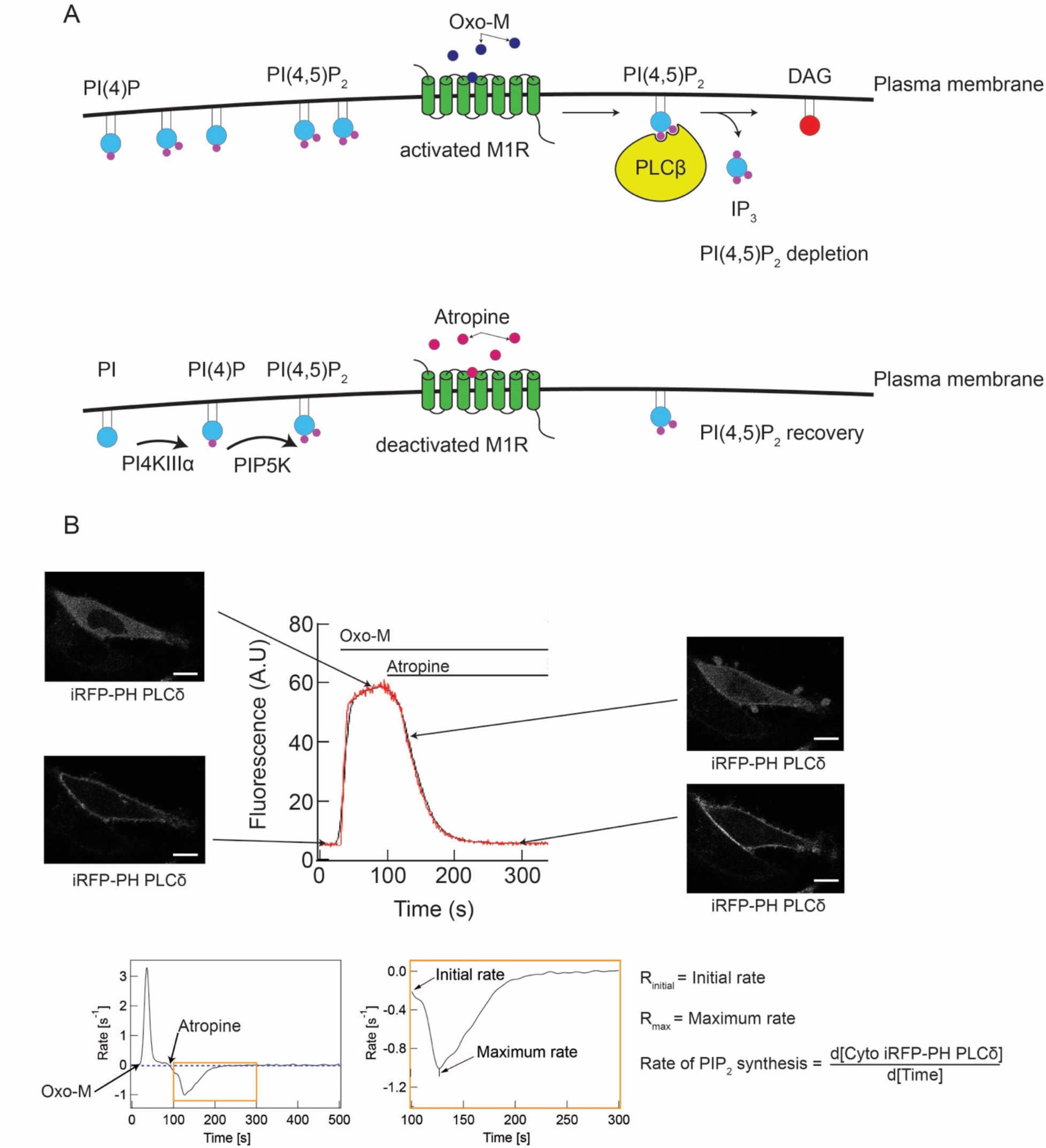
Experimental setup for studies to quantify kinetics of PI(4,5)P_2_ recovery after acute depletion. (A) Diagram representing the M1R-mediated PI(4,5)P_2_ depletion system. Oxo-M, an agonist of M1R, is used to activate the receptor, resulting in downstream activation of PLCβ and PI(4,5)P_2_ depletion at the PM. Addition of atropine, an antagonist of M1R, results in inactivation of PLCβ and recovery of PI(4,5)P_2_ at the PM. (B) Representative curve (red line is raw data and black line is smoothened curve) depicting the cytosolic fluorescence of the PI(4,5)P_2_ biosensor, iRFP-PH(PLCδ), over time with corresponding confocal micrographs of a cell expressing iRFP-PH(PLCδ) at different timepoints in the process. Scale bars: 15 μm. The inset, boxed in orange, depicts the relevant time period, post-addition of atropine, for determination of relevant kinetics parameters, which are defined at right: initial rate of PI(4,5)P_2_ resynthesis and maximum rate of PI(4,5)P_2_ resynthesis.

We evaluated two kinetic parameters of PI(4,5)P_2_ resynthesis process: initial rate and maximum rate (**Fig. 4B**). Rates were measured as the slope of the tangent to the recovery curve at a specified time point. The initial rate corresponded to the rate of loss of the cytosolic fluorescent signal of iRFP-PH(PLCδ), at t=0 of the recovery (i.e., the addition of atropine). The maximum rate corresponded to the highest rate of loss of iRFP-PH(PLCδ) from the cytosol, i.e., the point of the recovery curve for which the slope of the tangent is highest. These parameters afforded a comprehensive view of the recovery of the PI(4,5)P_2_ biosensor, iRFP-PH(PLCδ), to the PM. Because the conversion of PI to PI(4)P is the rate-limiting step in the resynthesis of PI(4,5)P_2_^[39]^, monitoring PI(4,5)P_2_ resynthesis serves as a useful proxy for the PI 4-kinase activity in this system.

We began by examining the initial rates of PI(4,5)P_2_ resynthesis in control cells or cells overexpressing TMEM150A and/or EFR3B (either WT or a dual-palm mutant). We found that overexpression of TMEM150A did not result in a statistically significant increase in the initial rate of PI(4,5)P_2_ synthesis, though it showed a trend in that direction, in general agreement with published data^[20]^, (**Fig. 5A**). Compared to TMEM150A, however, overexpression of EFR3B led to a much larger increase in the initial rate of PI(4,5)P_2_ resynthesis. Though none of the dual-palm mutants had the same effect as WT EFR3B, overexpression of the C7S and C8S mutants resulted in a small but significant increase in the measured initial rate, consistent with a role in the formation of Complex I. Furthermore, we found that co-expression of TMEM150A with WT EFR3B or the C7S and C8S EFR3B mutants had no effect on the initial rate of PI(4,5)P_2_ synthesis (**Fig. 5B**). However, co-expression of the EFR3B(C5S) mutant with TMEM150A had a stronger effect on the initial rate than expression of the C5S mutant or TMEM150A alone. Based on these comparisons, we conclude that PI 4-kinase activity from Complex I impacts the initial rate more strongly than that from Complex II, which may form on a slower timescale following atropine treatment. However, the co-overexpression of the EFR3B(C5S) mutant, which preferentially interacts with TMEM150A (see iDRM results from **Fig. 3**), may generate Complex II on a faster timescale, resulting in a statistically significant increase in the initial rate of PI(4,5)P_2_ resynthesis (**Fig. 5B**).

**Figure 5.**
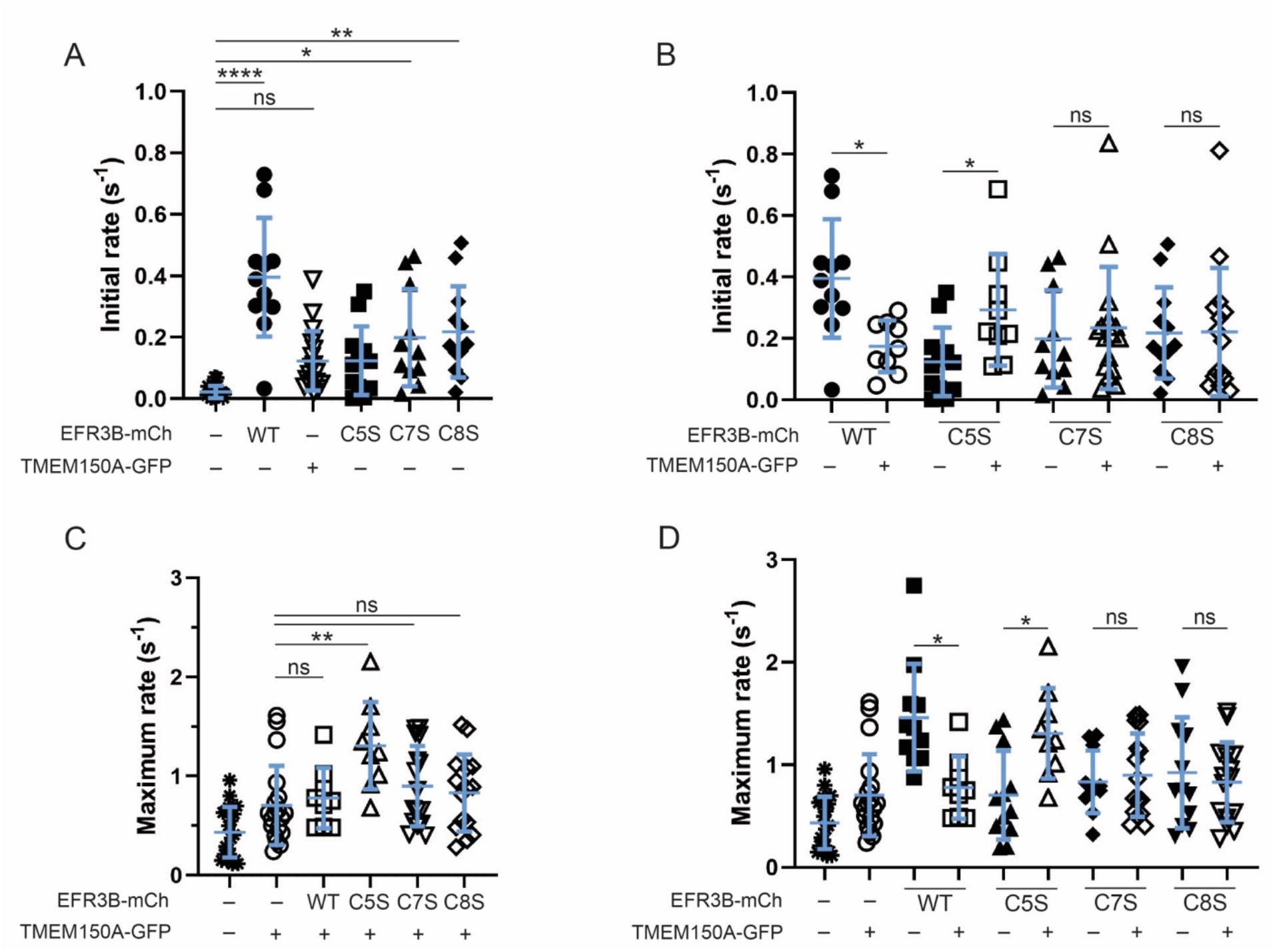
The C5S mutant of EFR3B facilitates a more rapid recovery of PI(4,5)P_2_ homeostasis in combination with TMEM150A than other palmitoylated forms of EFR3B. HeLa cells were transfected with iRFP-PH(PLCδ), either M1R-YFP or M1R-3xFLAG, and the indicated combination of TMEM150A-GFP and/or an EFR3B-mCherry variant (WT or CxS mutant). A time-lapse movie was acquired, recording iRFP fluorescence every 1.2 s, for 10 min. At t=30 s, oxo-M was added to induce PI(4,5)P_2_ hydrolysis, and at t=90 s, atropine was added to stop PI(4,5)P_2_ hydrolysis, permitting its resynthesis from PI, via PI(4)P. Shown are kinetics parameters during the post-atropine phase, as described in the legend of **Fig. 4**: initial rate of PI(4,5)P_2_ recovery (**A** and **B**) and maximum rate of PI(4,5)P_2_ recovery (**C** and **D**). Note that, to improve clarity and facilitate comparisons between effects of either EFR3B mutants alone or such mutants when co-expressed with TMEM150A, some data in (**A**) also appears in (**B**); similarly, some data in (**C**) also appears in (**D**). Statistical significance was determined using the one-way ANOVA with Tukey-Kramer post-hoc test. n=9–21. *, p<0.05; **, p<0.025; ***, p<0.01; ****; p<0.001; ns, not significant.

We next investigated additional kinetic parameters of the recovery of PM pool of the PI(4,5)P_2_ biosensor to glean insight into potential effects on both Complexes I and II. First, we found that the maximum rate of PI(4,5)P_2_ recovery was strongly increased in cells expressing WT EFR3B and more modestly increased upon overexpression of any of the dual-palm mutants (**Fig. 5D**, filled shapes). Second, overexpression of TMEM150A modestly increased the maximum rate of PI(4,5)P_2_ recovery compared to control cells (**Fig. 5C**, hollow vs. filled circles). Compared to this modest increase, systematic co-overexpression of TMEM150A with either WT EFR3B or each of the dual-palm mutants revealed that only C5S co-overexpression led to a further increase in maximum rate of PI(4,5)P_2_ recovery (**Fig. 5C–D**). Overall, these observations support the idea that the EFR3B(C5S)- and TMEM150A-containing Complex II is most efficient at promoting PI(4)P production for PI(4,5)P_2_ resynthesis after acute depletion and strengthens the implication of TMEM150A in PI(4,5)P_2_ homeostasis.

## DISCUSSION

PI4KIIIα, the enzyme responsible for the majority of PI(4)P synthesis at the PM, is thought to exist in two distinct complexes, containing either EFR3/TTC7/FAM126 (Complex I) or EFR3/TMEM150A (Complex II) (**Fig. 1A**)^[20]^. The goals of this study were to elucidate mechanisms underlying the partitioning of PI4KIIIα between these two complexes and reveal functional differences between them in support of PI(4)P synthesis at the PM. Because the sole shared component between these two PI4KIIIα complexes is EFR3, whose membrane anchoring requires a Cys-rich palmitoylation motif, we posited that differential localization within Lo-like and Ld-like regions of the PM may be a defining difference between Complexes I and II critical to their unique functional properties.

PI(4,5)P_2_ is enriched in Lo-like regions of the PM^[28,40–42]^, and some evidence points to its local synthesis in these membrane domains^[28,43]^. However, several studies have also detected substantial pools of PI(4,5)P_2_ in Ld-like membrane domains^[28,40,42]^. The mechanisms that lead to specific synthesis and enrichment of PI(4,5)P_2_ in Lo or Ld domains are not well understood. Clearly, multiple lines of evidence point to a primary role of PI4KIIIα in the synthesis of PI(4,5)P_2_, via production of PI(4)P, in the PM^[8,44]^. The link between TMEM150A and PI(4,5)P_2_ resynthesis following acute depletion may shed some light on the compartmentalization of PI(4,5)P_2_ in the PM. Specifically, the capacity of EFR3 for clustered palmitoylation at its N terminus and the well-established connection between palmitoylation and Lo partitioning of proteins^[27]^ prompted us to investigate the membrane partitioning properties of EFR3 and their impact on its interaction with TMEM150A. We addressed the above questions using a multidisciplinary approach involving chemical biology, biochemistry, and fluorescence imaging techniques. Our results provide evidence for a palmitoylation code on EFR3B, the membrane anchor of both PI4KIIIα complexes, which drives the selective formation of the TMEM150A-containing Complex II (**Fig. 6**).

**Figure 6:**
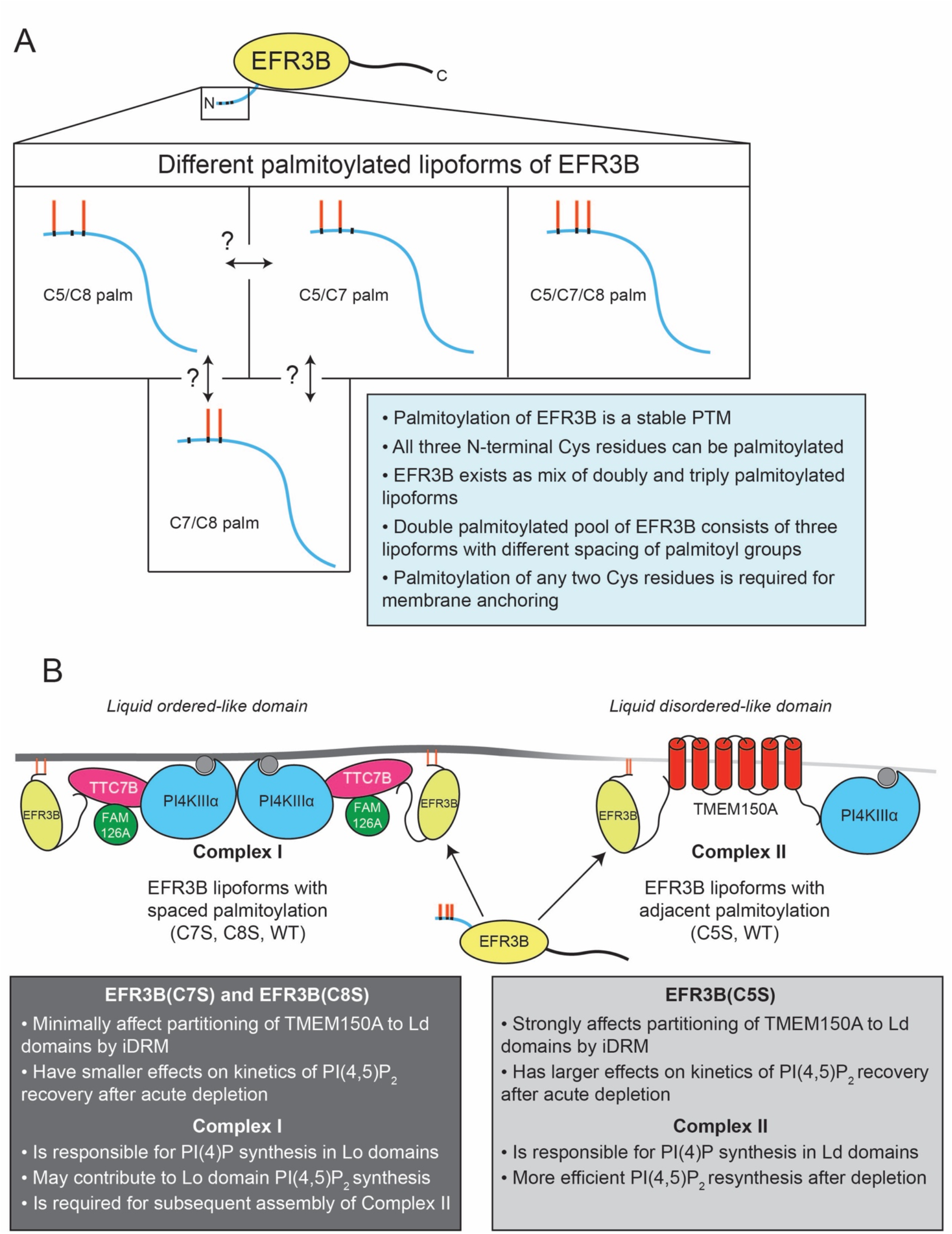
A model for the regulation of the formation of the TMEM150A PI4KIIIα complex. A) Schematic representation of the N-terminus of EFR3B and its multiple membrane-anchored dual and triply palmitoylated lipoforms. The C7S and C8S mutants preferentially form Complex I, whereas the C5S mutant interacts with TMEM150A to support assembly of Complex II. Whether lipoforms of EFR3B can be interconverted from one to the other is unknown. B) Proposed model for the regulation of the formation of the two distinct PI4KIIIα complexes. Functional studies indicate that conditions favoring formation of Complex II results in higher PI4KIIIα activity, positively impacting PI(4,5)P_2_ homeostasis.

Using acyl-PEG exchange, we found that, among the three potentially palmitoylated Cys residues in the EFR3B N-terminal motif, two or three are occupied in the WT protein and mutation of any one Cys resulted in near-quantitative palmitoylation of the remaining two palmitoylation sites. These results show that all three Cys residues can be palmitoylated in live cells and that palmitoylation of any two is sufficient for PM localization of EFR3B. Using pulse-chase metabolic labeling with a bioorthogonal palmitate analog, we further showed that the palmitoylation of EFR3B is highly stable, allowing for robust, long-term PM anchoring. Collectively, these results suggest that the dual palmitoylated pool of WT EFR3B observed by acyl-PEG exchange is likely a heterogenous mix of the three palmitoylation combinations at C5/C7, C5/C8, and C7/C8, giving rise to an ensemble of palmitoylation patterns on EFR3B. We note that the pulse-chase experiments do not preclude the enzymatic or non-enzymatic migration of palmitoyl groups within this motif in the dual palmitoylated forms of EFR3B.

The palmitoylation status of EFR3B affected its biophysical organization within the PM. Palmitoylation of peripheral membrane proteins is a primary regulator of their preference for Lo-like nanodomains^[27]^. Indeed, our quantitative iDRM results reveal that WT EFR3B is moderately detergent-resistant, supporting a Lo-like domain preference. However, its Lo-preference was not as strong as other well-known Lo-preferring lipidated protein probes, e.g., Lyn kinase ^[33]^. The detergent resistance of WT EFR3B (consisting of both triply and doubly palmitoylated species) was very similar those of the dual palmitoylated CxS mutants, indicating that the triply palmitoylated WT EFR3B species had a similar Lo-preference to its doubly palmitoylated counterparts. This similarity is also reflected in the similar diffusion speeds (*t*_1/2_ values in FRAP experiments) of WT and all three CxS EFR3B mutants. The moderate Lo-preference of EFR3B also indicates that a substantial minority fraction of EFR3B is present in Ld-like domains at steadystate. Having pools of EFR3, the membrane anchor for all known PI4KIIIα complexes, in both Lo and Ld-pools could be functionally relevant, because PI(4,5)P_2_ is likely synthesized in both domains^[28,40,42]^, requiring recruitment of PI4KIIIα to both Lo- and Ld-like domains.

Our iDRM results demonstrated that TMEM150A is moderately resistant to detergent, indicating its Lo-preference. Transmembrane proteins had been previously thought to be excluded from the Lo-like domains due to unfavorable energetics^[45]^. However, some recent examples of Lo-preferring TM proteins have been reported, in which the biochemical basis of the Lo preference depends on the protein itself and its interaction with membrane lipids^[46–49]^. Molecular dynamics simulations using realistic lipid bilayer models revealed that membrane proteins can organize the lipid environment around themselves to maximize favorable interactions, further suggesting a preference for certain lipids^[50]^. TMEM150A is likely a new addition to this category of membrane proteins, although the exact physicochemical principles governing its Lo-like preference remain unknown. A recent study demonstrated that Sfk1, the yeast homolog of TMEM150A, acts to retain ergosterol, the yeast cholesterol analogue, in the PM^[51]^. This property may translate to TMEM150A in mammalian cells, where cholesterol is a key regulator of Lo-domain formation^[52]^. Interestingly, the detergent resistance of TMEM150A is highly sensitive to the presence of WT EFR3B or the C5S mutant, suggesting that the Lo partitioning of TMEM150A, or its proteinprotein interactions, are modulated by interactions with EFR3B, bringing it to Ld domains. The EFR3B C7S mutant showed no effect on TMEM150A partitioning, and the C8S mutant showed an intermediate effect. These results led us to hypothesize that the adjacent palmitoylation sites on the EFR3B C5S mutant leads to favorable interactions with TMEM150A, thus driving the formation of the Complex II in Ld-like membrane regions.

Interestingly, we observed an increase in the detergent resistance of WT EFR3B (consisting of both triply and doubly palmitoylated species) in the presence of TMEM150A, whereas none of the dual-palm mutants showed this behavior. Separately, we confirmed that the relative abundance of the double and triple palmitoylated pools of WT EFR3B are unchanged by the presence of TMEM150A. Because the EFR3B CxS mutants did not exhibit any differences, relative to WT EFR3B, in detergent-resistant fraction by iDRM, diffusion kinetics and mobile fraction by FRAP, we hypothesize that the triply palmitoylated lipoform of EFR3B partitions more strongly into Lo-like domains in the presence of TMEM150A. This notion is consistent with the reduction in lateral diffusion of WT EFR3B measured by FRAP in the presence of TMEM150A. However, the underlying mechanism that confers stronger Lo-preference of the triply palmitoylated EFR3B only in the presence of TMEM150A is currently unknown.

The iDRM measurements on PI4KIIIα in the fully assembled Complex I revealed strong detergent resistance, suggesting an Lo-like membrane environment surrounding this complex. This observation is consistent with the inherent Lo-preference of EFR3B, the sole membrane anchoring component of this complex. The scenario is different for Complex II, which is formed by overexpression of TTC7B, EFR3B, TMEM150A and PI4KIIIα but only contains the latter three components^[20]^. Complex II has two constitutively PM-localized components, EFR3B and TMEM150A. Under these conditions of forced overexpression these four proteins (which likely results in formation of a mixture of complexes I and II), we observed that PI4KIIIα is resistant to detergent but TMEM150A is not, meaning that the TMEM150A-containing complex is Ld-preferring. The *R* values from iDRM experiments of PI4KIIIα under conditions favoring formation of Complex I and Complex II are strikingly similar. These observations suggest that PI4KIIIα does not exist exclusively in one complex or the other when all four components are co-transfected but rather that the two complexes likely co-exist in the PM, each with different preferences for Lo-like (Complex I) and Ld-like membrane domains (Complex II) at steady state (**Fig. 6**).

In addition to revealing the biophysical organization of PI4KIIIα at the PM, our data also support a functional role for an EFR3B palmitoylation code on the activity of PI4KIIIα in the context of PI(4,5)P_2_ resynthesis after acute depletion. Among the three EFR3B dual-palm mutants, only C5S interacts favorably with TMEM150A, resulting in more efficient resynthesis of PI(4,5)P_2_ after acute depletion. Our data are consistent with a known role for TMEM150A in PI(4,5)P_2_ homeostasis^[20]^. We show that, upon co-expression of EFR3B(C5S) and TMEM150A, cells exhibit a faster maximum rate of PI(4,5)P_2_ resynthesis compared to cells expressing TMEM150A with the either EFR3B(C7S) or EFR3B(C8S). Therefore, the specific interaction of TMEM150A with the EFR3B(C5S) mutant in Ld-like domains, but not with other EFR3B dual-palm mutants, has the strongest effect on PI(4,5)P_2_ production after acute hydrolysis. Importantly, the use of PI(4,5)P_2_ recovery experiments here is supported by previous findings indicating that phosphorylation of PI to PI(4)P, rather than the subsequent conversion of PI(4)P to PI(4,5)P_2_, is the rate-limiting step in PI(4,5)P_2_ resynthesis following agonist-mediated M1R stimulation^[39,53]^.

These results support a model wherein Complex II is in part responsible for the synthesis of PI(4,5)P_2_ in Ld-like membrane regions, whereas Complex I partitions to Lo-like membrane regions, where the PI(4)P it produces may be used to replenish PI(4,5)P_2_ stores in these membrane domains. We propose that an equilibrium exists between the two PI4KIIIα-containing complexes that is in part dictated by the relative abundance of the different lipoforms of EFR3B, with the lipoform containing adjacent palmitoyl groups at C7/C8 preferentially forming Complex II with TMEM150A, and the other two lipoforms with palmitoyl groups spaced farther apart preferentially forming Complex I. Considering the well-known preference of palmitoylation for Lo-like membrane domains^[27]^, it is likely that Complex I shows a preference for Lo-like membrane domains. Relative to Complex I, Complex II shows a decreased preference for Lo-like nanodomains and therefore an increased affinity for Ld-like nanodomains. PI(4,5)P_2_ pools in Lo and Ld domains, synthesized by different PI4P 5-kinase isoforms, play different functional roles in calcium signaling^[42,43]^. Specifically, Ld pools of PI(4,5)P_2_ reduce store-operated calcium entry (SOCE)^[42]^. In light of these findings, our results raise the possibility that the Complex II may contribute to PI(4,5)P_2_ synthesis in Ld domains to regulate calcium signaling in response to SOCE, which is activated by GPCRs that signal via Gq, such as M1R.

The idea of Complex I favoring Lo-like domains has support by several lines of indirect evidence from the literature. First, yeast Stt4 exists at the PM in so-called PIK patches with the EFR3 and TTC7 orthologs, Efr3 and Ypp1, that contain ~30 copies of each protein, and which recover minimally from photobleaching on the minute timescale^[19]^. Second, the plant orthologs of EFR3, TTC7, and FAM126, were recently identified, and their roles in recruiting the plant PI4KIIIα ortholog, PI4Kα1, to the PM were conserved. Interestingly, the EFR3 ortholog EFOP (EFR3 of Plants) is PM-localized via palmitoylation of multiple Cys residues, and the complex appears to associate with specific membrane nanodomains^[54]^.

Many questions remain about the function of TMEM150A and its interactions with EFR3. The determinants of the protein-protein interaction between TMEM150A and EFR3B, including how the former can distinguish different EFR3B lipoforms, are unknown, although the C-terminal cytosolic tail of TMEM150A has been implicated in this interaction and in its role PI(4,5)P_2_ homeostasis^[20]^. Furthermore, the palmitoylation state of EFR3A, which contains four adjacent Cys residues within its N-terminal palmitoylation motif, is not known, and would be an interesting area for future study. Moreover, further experiments are needed to establish the relative abundance of each lipoform of EFR3B at endogenous expression levels. In yeast, Efr3 is not palmitoylated^[8]^ and exclusively relies on its polybasic patch to interact with the PM, highlighting a major mechanistic difference in the assembly of PI4KIIIα complex in this organism. Though Sfk1, the yeast homolog of TMEM150A, is clearly involved kinase recruitment and function^[18]^, the mechanism by which it does so, and its relationship to the Efr3–Ypp1–Stt4 complex, remains unknown. The roles of Sfk1 in retention of ergosterol in the PM^[51]^ and maintenance of PM integrity and impermeability^[55]^ may or may not be related to its role in Stt4 function. Reciprocally, it is unknown if TMEM150A plays a similar role in PM cholesterol homeostasis, transbilayer movement of lipids, or membrane integrity in mammalian cells.

In conclusion, we propose that a palmitoylation code within EFR3B, the membrane anchor for PI4KIIIα in the PM, dictates the partitioning of PI4KIIIα between two distinct multicomponent complexes responsible for production of PI(4)P in this membrane. Our model suggests unique roles for distinct lipoforms of EFR3B with very subtle differences in their palmitoylation patterns, in influencing interactions with TMEM150A and localization of PI4KIIIα to Lo-like and Ld-like domains. We show that these subtle changes in EFR3B palmitoylation impact the function of PI4KIIIα in producing PI(4)P required for restoration of PI(4,5)P_2_ homeostasis at the PM following its acute depletion by Gq–PLC signaling. These findings reinforce the value of evaluating the detailed mechanisms of posttranslational lipidation in the spatiotemporal regulation of protein–protein and protein–lipid interactions within the PM, and we anticipate that similar such functionally important palmitoylation codes on multiply palmitoylated proteins will emerge in diverse contexts.

## AUTHOR CONTRIBUTIONS

A.G.B. performed confocal microscopy, biochemical APE and alk-16 assays, and molecular cloning, N.B. and A.G.B. performed FRAP and PI(4,5)P_2_ recovery experiments, N.B. performed ImFCS experiments, and N.B. and H.P. performed iDRM experiments. A.G.B., N.B., B.A.B., and J.M.B. designed experiments and interpreted data. A.G.B., N.B., and J.M.B. wrote the manuscript, with edits and input received from all other authors.

## ACKNOWLEDGMENTS

J.M.B. acknowledges support from the NIH (R00GM110121 and R01GM131101) and the Alfred P. Sloan Foundation (Sloan Research Fellowship). B.A.B. acknowledges support from NIH (R01GM117552). We thank Chris Diaz for technical assistance with cloning, Allan Lee for help performing DRM experiments, and Alice Wagenknecht-Wiesner for her help in maintaining cells and plasmids. We thank the Fromme and Emr labs for sharing equipment, and David Holowka, Maurine Linder, Scott Emr, and members of the Baskin and Baird-Holowka labs for helpful discussions.

## MATERIALS AND METHODS

### General Methods

Sources of chemical reagents, primers, antibodies, and plasmids are listed in **Table S1**.

### Cell culture

All cell lines were cultured at 37 °C in 5% CO_2_. HeLa cells were grown in DMEM (Corning) supplemented with 10% FBS and 1% penicillin/streptomycin (P/S). RBL-2H3 cells were maintained in MEM (Corning) supplemented with 20% FBS and 10 mg/L gentamicin sulfate.

### Transfection

#### Chemical transfection of HeLa cells

Cells were seeded on #1.5 glass-bottom, 35 mm imaging dishes (MatTek and Matsunami) at a density of 150,000 cells per dish and left to grow overnight. Transfections were carried out the next day. In a polystyrene tube, 150 μL of Transfectagro (Corning) were added per transfected dish and 1.5 μL of Lipofectamine 2000 (Thermo Fisher) per dish were mixed in. In a separate transfection tube, the same volume of Transfectagro was added and mixed with 1.5 μg of each transfected plasmid. Tubes were left to incubate for 5 min at room temperature and then mixed together. After mixing by gentle pipetting, tubes were left at room temperature for 20 min. The media from the imaging dishes was aspirated and replaced with 1.7 mL of Transfectagro supplemented with 10% FBS. 300 μL of the appropriate transfection mix was added to each dish. Dishes were placed in an incubator at 37 °C. After 7 h, Transfectagro was aspirated and replaced with growth medium, and the cells were left in the incubator for imaging the next day.

The transfection procedure for 60 mm dishes is the same except that dishes are seeded with 800,000 cells, and 375 μL of Transfectagro were used, for a total of 750 μL per dish. 4.5 μg of each plasmid were used for each dish and 4 μL of Lipofectamine 2000 were used for each dish.

#### Chemical transfection of RBL-2H3 cells

20,000 cells were suspended in 2 mL of growth medium and placed, homogeneously spread, in a 35 mm glass-bottom imaging dish (MatTek). After overnight growth, the cells were transfected using FuGENE HD transfection kit (Promega). For one imaging dish, plasmid DNA (2 μg of each plasmid) and FuGENE (3 μL FuGENE/μg DNA) were mixed thoroughly in 100 μL Opti-MEM (Thermo Fisher) and incubated at room temperature for 15 min. Next, cells were washed once and covered with 1 mL Opti-MEM. The DNA/FuGENE complex was spread evenly over the cells and incubated for 1 h. 1 mL of prewarmed phorbol 12,13-dibutyrate (PDB, 0.1 μg/mL; diluted in Opti-MEM from a 10,000X stock prepared in DMSO) was then added to each imaging dish and cells were incubated for 3 h at 37 °C. Finally, 2 mL of growth medium was added to each imaging dish after discarding Opti-MEM. The transfected cells were cultured for 18–22 h before FRAP measurements.

#### Electroporation of RBL-2H3 cells

RBL-2H3 cells in a confluent 75 cm^2^ flask were washed and trypsinized for 8 min at 37 °C with 3 mL of 0.05% Trypsin-EDTA (Thermo Fisher). The detached cells were resuspended in 7 mL of growth medium and centrifuged to remove the medium. The cell pellet (1.5 x 10^6^ cells) was resuspended in 1.5 mL of cold electroporation buffer (137 mM NaCl, 2.7 mM KCl, 1.0 mM MgCl_2_, 1 mg/mL glucose, and 20 mM HEPES; pH 7.4). Next, 5 μg of plasmid DNA corresponding to each protein of interest was thoroughly mixed with 500 μL of the resuspended cells in an electroporation cuvette (Bio-Rad). This cuvette was subject to an electroporation pulse (280 V, 950 μF) using a Gene Pulser X (Bio-Rad) electroporation module. The electroporated cells were then added to 6 mL of growth medium, mixed thoroughly, and deposited in imaging dishes (2 mL/dish). The cells were allowed to attach on the dish for 3 h at 37 °C, following which the medium was replaced with fresh growth medium. The cells were cultured for 24 h to recover before proceeding to the next sample preparation steps for iDRM.

### Cloning

WT EFR3B-3xFLAG, EFR3B(C5,7,8S)-GFP (mouse)^[8]^ and TMEM150A-GFP (human)^[20]^ are identical to constructs used in the referenced studies^[20]^, which were cloned in the p3xFLAG-CMV-14 and pEGFP-N1 vectors, respectively. EFR3B CxS and CxxS mutants were generated by using Quikchange PCR with the primers listed in **Table S1** and validated by sequencing. C7S, C8S and C7,8S mutants were generated using WT EFR3B as a template. C5S, C5,7S and C5,8S mutants were generated using EFR3B(C5,7,8S) as a template. M1R-3xFLAG was subcloned from M1R-YFP, using NotI and EcoRI. Subcloning was performed using appropriate restriction enzyme pairs compatible with donor and target plasmids according to standard procedures.

### Acyl-PEG exchange palmitoylation assay

Acyl-PEG exchange experiments were performed essentially as previously reported^[29]^. Briefly, cells were lysed in lysis buffer (50 mM triethanolamine (TEA), 4% SDS, 150 NaCl, pH 7.3 with 1x cOmplete Protease Inhibitor cocktail (Roche)). Cell lysates were sonicated to ensure complete lysis and to fragment nucleic acids. Protein concentration was determined by BCA assay and protein concentration was normalized to 2 mg/mL. For the assay, 92.5 μL of protein were used. 5 μL of 200 mM TCEP (neutralized to pH 7 by addition of 800 mM NaOH) were added to all samples, which were then left on the nutator at room temperature for 30 min. 2.5 μL of 1 M N-ethylmaleimide (NEM) in ethanol were added to all samples. Samples were then incubated for 2 h at room temperature on the nutator. A protein precipitation was then performed by adding prechilled methanol, water and chloroform at a 400 μL:150 μL:300 μL ratio. Samples were inverted and spun at 17,000 x g for 5 min at 4 °C. The aqueous layer (top) was gently removed and replaced with 1 mL of methanol. Samples were spun again in the same conditions for 3 min, and the supernatant removed and replaced with 800 μL of methanol. Samples were spun again for 3 min. The methanol was decanted, and samples dried on a benchtop concentrator (CentriVap) at 37 °C until no methanol remained. Samples were resuspended in 100 μL of lysis buffer in a 37 °C water bath for 10 min, or until protein pellet was no longer visible. Samples were then gently sonicated in a sonicating water bath for 10 s to ensure total dissolution of the pellet. This protein extraction was performed a total of three times. After the third time, samples were resuspended in 60 μL of lysis buffer with EDTA and without protease inhibitor. After pellet dissolution, 90 μL of 1 M hydroxylamine (pH 7) dissolved in 50 mM TEA 0.2% TX-100 were added to each sample. For a no hydroxylamine control, buffer with 50 mM TEA 0.2% TX-100 was used. Samples were placed on the nutator for 1 h at room temperature, and then a protein precipitation was performed, after which pellets were solubilized in 30 μL of lysis buffer and treated with 90 μL of a solution of 1.33 mM mPEG-maleimide in 50 mM TEA and 0.2% TX-100. Samples were left on the nutator at room temperature for 2 h. A final protein precipitation was performed, after which samples were resuspended in 50 μL of lysis buffer and resolubilized. SDS-PAGE loading buffer (6X Laemmli) was added, and samples boiled at 95 °C for 5 min and stored at −20 °C until analysis by SDS-PAGE/Western blot as described below.

### Bioorthogonal metabolic labeling pulse-chase assay for palmitoylation stability

HeLa cells were seeded on 60 mm dishes (800,000 cells per dish). The next day, cells were transfected according to the above protocol with EFR3B-3xFLAG. After 24 h, cells were rinsed with PBS, and the growth medium changed to DMEM supplemented with 10% dialyzed FBS containing 50 μM 17-ODYA (alk-16). The next day, the 17-ODYA was rinsed out of the dishes and the growth medium replaced with regular growth media (i.e., DMEM with standard FBS and P/S) in the presence of lactacystin (1 μM) and cycloheximide (50 μg/mL) for 1, 2, 4, or 8 h. After each time point, cells were harvested, rinsed three times in PBS by pelleting at 4 °C in a benchtop centrifuge (900 x g, 2 min) in between washes. Cells were either flash frozen in liquid nitrogen for temporary storage or lysed in 200 μL of lysis buffer (50 mM TEA, pH 7.5, 150 NaCl, 1% NP-40, 0.25% sodium deoxycholate, and cOmplete EDTA-free protease inhibitor cocktail). Samples were then sonicated while taking care to keep them on ice. Protein concentration was normalized to 2 mg/mL by performing a BCA assay. An anti-FLAG IP to recover labelled EFR3B was performed using 500 μg of sample and 20 μL of anti-FLAG bead slurry (EZview Red ANTI-FLAG M2 Affinity gel from Millipore Sigma). Samples were left to rotate for 2 h at 4 °C in a cold room. The beads were then recovered and rinsed three times with wash buffer (25 mM TEA, 150 mM NaCl, 0.2% NP-40, pH 7.4) with pelleting in between each step (900 x g, 2 min). After the last rinse, beads were resuspended in 16 μL of the wash buffer. Cy5.5-azide was then tagged onto the 17-ODYA by the Cu-catalyzed azide-alkyne cycloaddition (click chemistry). The reaction requires final concentrations of 100 μM Cy5.5-azide, 1 mM CuSO_4_, 600 μM THPTA and 50 mM sodium ascorbate. 20X stocks were made in DMSO for Cy5.5-azide and in water for CuSO_4_, THPTA, and sodium ascorbate. The CuSO_4_ and sodium ascorbate solutions were made fresh every time. To carry out the reaction, the CuSO_4_ and THPTA solutions were first mixed in a 1:1 ratio. Then, 1 μL of the Cy5.5-azide solution was added to the samples, followed by 2 μL of the pre-mixed CuSO_4_/THPTA solution, and finally 1 μL of the sodium ascorbate solution. The reactions were left at room temperature in the dark for 1 h. After the reaction was completed, beads were rinsed three times with the wash buffer to remove excess fluorophore. Samples were then denatured using 6X SDS-PAGE Laemmli loading buffer diluted down to 2X. Samples were boiled at 100 °C for 5 min and stored at −20 °C until analysis by SDS-PAGE/Western blot as described below.

### SDS-PAGE and Western blot

SDS-PAGE was performed using the Mini-Protean system (Bio-Rad), running at 160 V using Precision Plus molecular weight standards (Bio-Rad). For Western blot, transfers to nitrocellulose were carried out for 2 h at 70 V (constant voltage) at 4 °C. Membranes were stained using Ponceau S and scanned on an image (ChemiDoc MP, Bio-Rad). Blocking was performed for 1 h with rocking at room temperature using 5% milk in TBS + 0.1% Tween-20 (TBS-T). Primary antibody staining was performed overnight at 4 °C with rocking, with primary antibody diluted in blocking buffer. Following 3 x 5 min rinses in TBS-T, secondary antibody incubation was performed with the appropriate secondary antibody-HRP fusion (1:5,000 dilution in blocking buffer) for 1 h at room temperature with rocking. The membranes were then rinsed 3 x 7 min in TBS-T, and 7 min in TBS. Membranes were then exposed to Clarity ECL solution (Bio-Rad) for 5 min and then imaged (Chemidoc MP, Bio-Rad). Western blots were quantified using the ImageLab software (Bio-Rad).

### Imaging detergent resistant membrane (iDRM) experiments

#### Sample preparation

All iDRM preparations were done on RBL-2H3 cells electroporated with a given set of plasmids and recovered overnight under physiological conditions. Cells in imaging dishes (MatTek) were washed with 1 mL of buffered salt solution (BSS: 135 mM NaCl, 5.0 mM KCl, 1.8 mM CaCl_2_, 1.0 mM MgCl_2_, 5.6 mM glucose, and 20 mM HEPES, pH 7.4) followed by addition of another 1 mL of fresh BSS. The dishes were placed on ice for 10 min. From the dishes designated as with TX-100 (+TX-100 sample), the BSS was removed slowly, and 1 mL of BSS with 0.04% TX-100 was added dropwise and incubated for 10 min in ice. A control dish (−TX-100) was prepared in which 1 mL of BSS without TX-100 was added dropwise for 10 min in ice. After this incubation step, the solutions were gently removed from both dishes to minimize any mechanical perturbation of the cells. 1 mL of fixing solution containing 4% paraformaldehyde and 0.1% glutaraldehyde in PBS was added dropwise to both dishes at room temperature for 20 min. The fixation was quenched by addition of 0.5 mL of 10 mg/mL bovine serum albumin (BSA) in PBS for 10 min, and samples were stored at 4 °C until imaging.

#### Imaging

iDRM was imaged with a home-built total internal reflection fluorescence microscope (TIRFM) involved using a home-built microscope (DMIRB, Leica Microsystems) with an oil immersion objective (PlanApo, 100X, NA 1.47; Leica Microsystems) and an electron-multiplying charge-coupled device camera (black illuminated Andor iXON 897DU, pixel size 16 μm, Andor Technology). To measure the fluorescence of proteins tagged with green fluorescent protein (GFP), a 488 nm laser (Coherent) was used. To measure the fluorescence of proteins tagged with mCherry, a 561 nm laser (Coherent) was used. Each image was taken with a data acquisition of 100 frames with 0.01 s per frame. Approximately 30 cells per dish were imaged. All images were saved in TIFF format in 16-bit grayscale with a data range of 0 to 65535. Image acquisition was performed with Andor Solis software. 30 cells each of the –TX-100 and +TX-100 samples for one biological replicate for a given set of probes. For each condition, at least three biological replicates were performed.

#### Data Analysis

Quantification and analysis were done using ImageJ/FIJI^[56]^. Cellular fluorescence was measured by manually drawing a region of interest (ROI) around the cell and using the measure feature of FIJI. To account for background fluorescence, a ROI was drawn outside of the cell, and the difference between the cell fluorescence and the background fluorescence was calculated to return a background-corrected average fluorescence. This process was performed for each cell in each construct with or without TX-100 samples. All images are the average of its 100 frames and are 200×200 pixels, in which pixel size is 160 nm. The background-corrected fluorescence of a probe in individual cells from –TX-100 and +TX-100 samples (~30 cells each) for given biological replicates yields a distribution of probe fluorescence values for each category. We used the Mann-Whitney U test to determine the statistical significance between –TX-100 and +TX-100 samples. Furthermore, we defined a parameter, called fluorescence retention (*R*) value, to quantitatively determine the extent of detergent resistance for a given probe using the following equation:

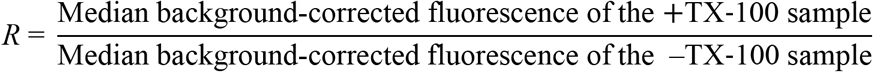

An *R* value of 1 means the probe is detergent-insoluble (i.e., the probe is surrounded by Lo-like environment), whereas a value of zero indicates complete solubility of the probe and its likely surrounding to be Ld-like. In practice, *R* values >0.6 are obtained for Lo-preferring lipid probes and <0.3 are obtained for Ld-preferring lipid probes^[33]^.

### Fluorescence recovery after photobleaching (FRAP) assays

Transfected cells were rinsed once with 1 mL of BSS to remove leftover growth medium. FRAP experiments were carried out in 1 mL of BSS buffer on a Zeiss LSM 800 confocal microscope equipped with a 1.4 NA, 40×, oil immersion objective and GaAsP detectors. Five frames were acquired before bleaching, after which cells were photobleached on their ventral surface using 488 nm laser illumination at 100% power for 50 iterations, resulting in strong visible bleaching in the target region of interest (bleached ROI). A separate, unbleached ROI of the same dimensions was used to control for photobleaching of the GFP tag due to image acquisition, and a third ROI was used to account for background fluorescence of the system (positioned outside of the cell). Time-lapse imaging was performed, with images acquired every 1.3 s for 3 min. All ROIs were circles of 3.5 μm diameter. FRAP data from the bleached ROI was normalized after accounting for photobleaching during recovery process and background using the fluorescence traces from control ROI (inside cell but away from bleached ROI) and background ROI (located outside cell) using the FRAPanalyzer software^[57]^. The normalized FRAP data were such that the fluorescence before photobleaching (i.e., F_normalized_ = 1 at recovery time (t) < 0) is 1 while at the time of bleaching is zero (i.e, F_normalized_ = 0 at t = 0). During and completion of the recovery the F_normalized_ is greater than zero (i.e., F_normalized_ >0 at t > 0). The normalized FRAP data (F_normalized_) as a function of recovery time (t) is then fitted using the following exponential function:

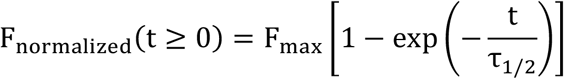

The saturation value of the fitted curve at extended time (i.e., when recovery is completed), F_max_, is the mobile fraction, whereas the timescale of diffusion is given by the recovery time (τ_1/2_) of the bleached spot.

### Imaging Fluorescence Correlation Spectroscopy (ImFCS)

The instrumental set-up, data acquisition and processing are described elsewhere^[35]^. Briefly, we collected time-lapse images (80,000 images at 3.5 ms frame rate) of ventral plasma membrane of fluorescently labelled RBL-2H3 cells using a home-built total internal reflection microscopy (TIRFM) equipped with high NA (1.49) oil-immersion objective and EMCCD camera (Andor iXon 897). Fluorescence fluctuations embedded in the image series was further processed by a FIJI/ImageJ^[56]^ macro (http://www.dbs.nus.edu.sg/lab/BFL/imfcs_image_j_plugin.html) to determine the *D* values.

### Imaging of PI(4,5)P_2_ resynthesis kinetics after acute depletion

#### Acquisition

HeLa cells were transfected with M1R-YFP or M1R-3xFLAG, iRFP-PH(PLCδ), and other constructs, as necessary. On the day of the experiment, imaging buffer (BSS + 1 mg/mL BSA) was warmed to 37 °C and used to rinse the cells once before the experiment. Confocal microscopy imaging was carried out on a Zeiss LSM 800 confocal microscope. Timelapse acquisitions were started prior to addition of oxo-M, with frames acquired every 1.2 s. After 30 s, oxo-M diluted in water was added to the cells at a final concentration of 10 μM. 60 s after the addition of oxo-M, atropine was added to the dish to a final concentration of 50 μM. The timelapse acquisition was continued for a total of 10 min. Recovery curves were fitted to an exponential using the Igor Pro software (Version 8; WaveMetrics). From the fitted curves, a value for *t*_1/2_ for recovery was obtained.

#### Data analysis

The time-dependent decrease of cytosolic fluorescence of iRFP-PH(PLCδ) was monitored after atropine addition. First, the fluorescence vs. time curve was smoothened to eliminate noise. The reliability of the smoothing process was confirmed by overlaying the smoothened curve on the raw curve. The PI(4,5)P_2_ synthesis rate at each point after agonist/antagonist addition was calculated from the derivative of smoothened fluorescence intensity with respect to time. Because we monitor cytosolic fluorescence, which decreases over time as PI(4,5)P_2_ is synthesized at the PM, the sign of the derivative values is negative. The derivative value at the first timepoint after atropine addition is termed as initial rate. The minimum of the derivative plot (i.e., the most negative value) corresponds to the maximum rate. The time difference between atropine addition and reaching the maximum rate is defined as time to reach max rate. Because the rate becomes zero at very long times after atropine addition, as no new net PI(4,5)P_2_ is being synthesized, this phenomenon is represented in the derivative plot as a plateau with average value of zero. All analysis was done in the Igor Pro software.

