## Supplementary Information for "A palmitoylation code controls PI4KIIIα complex formation and PI(4,5)P_2_ homeostasis at the plasma membrane"

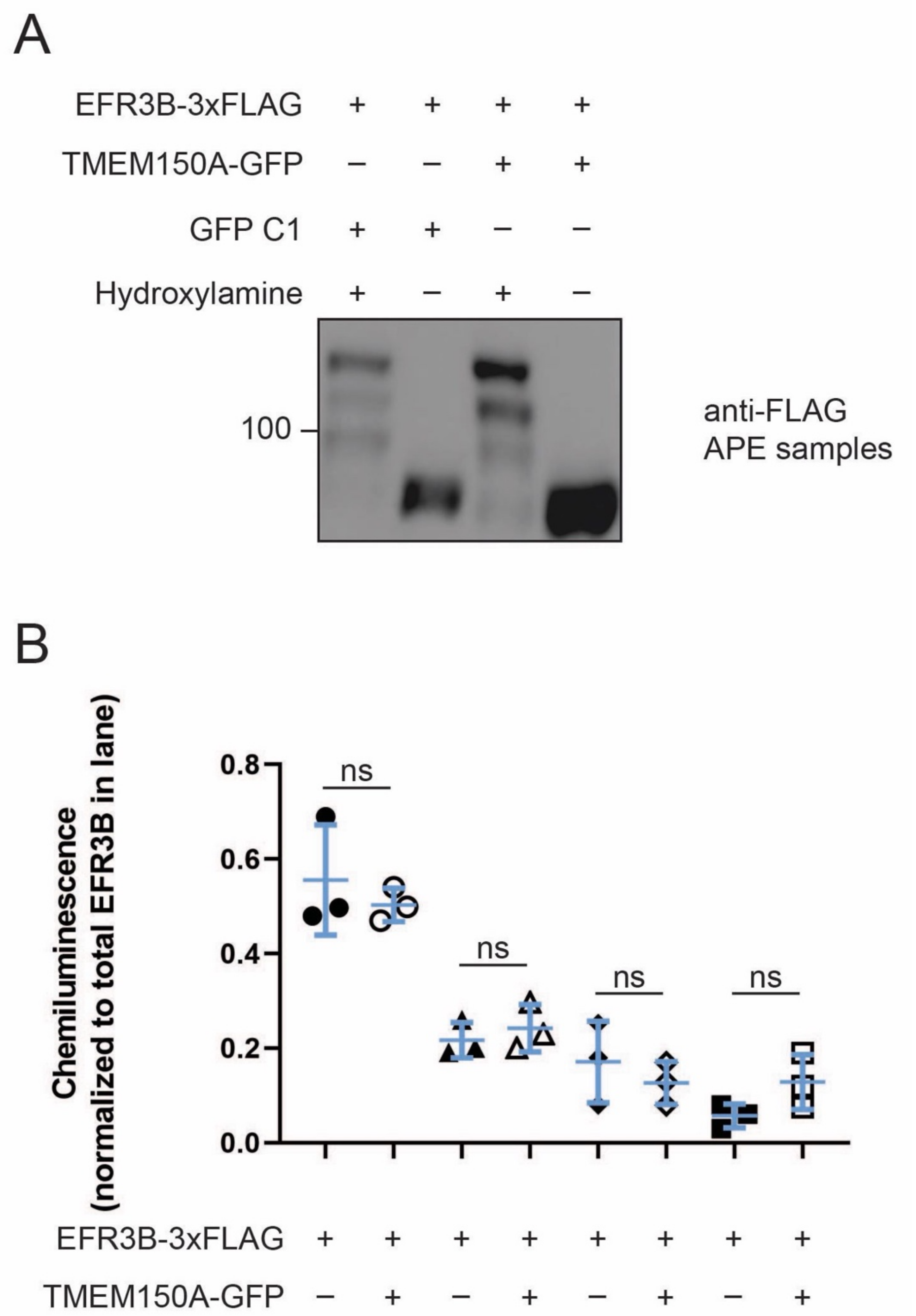

**Figure S1. Overexpression of TMEM150A does not alter the palmitoylation state of EFR3B.** (A) Representative Western blot of acyl-PEG exchange (APE) experiments in which EFR3B was co-expressed with TMEM150A-GFP or with a GFP-C1 empty vector. (B) Quantification of three biological replicates of the experiment shown in (A), showing that co-expression of TMEM150A with EFR3B has no effect on its palmitoylation. Statistical significance was assessed using Student's t-test. n=3; ns, not significant.

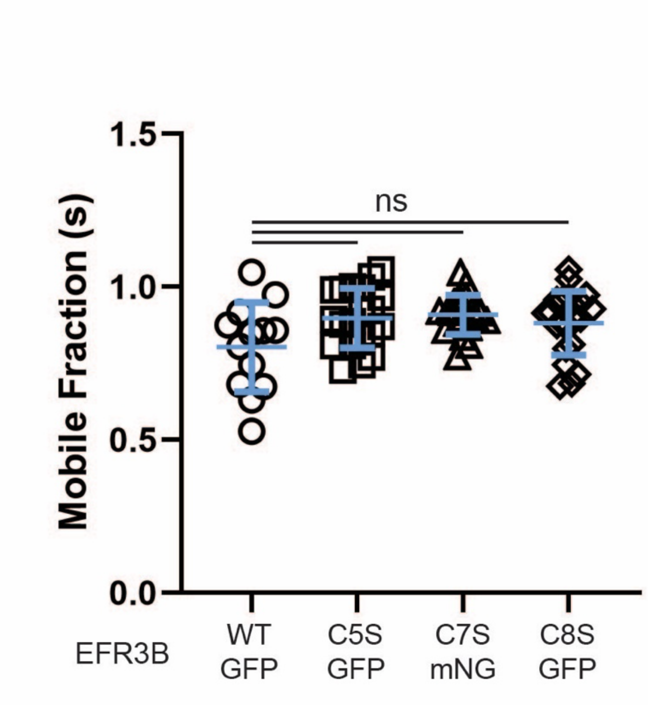

**Figure S2. The mobile fractions of WT and CxS EFR3B are the same.** Mobile fractions (mean ± SD) of the indicated EFR3B-GFP (or, for C7S, -mNG) construct, measured by FRAP. Note that EFR3B(C7S)-mNG was used due to undesired clustering of EFR3B(C7S)-GFP. n=13–22; ns, not significant.

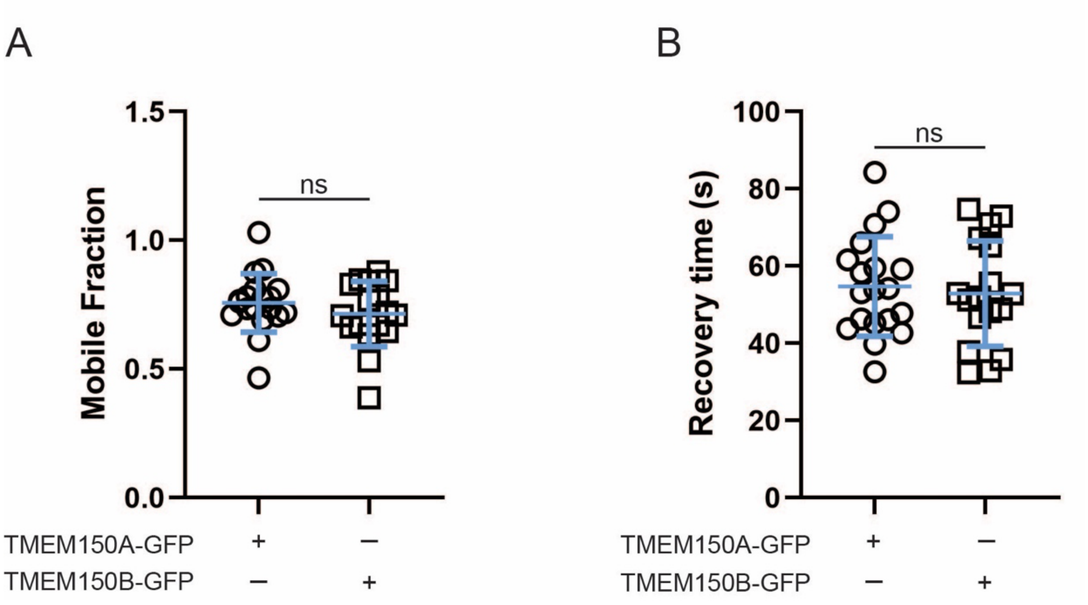

**Figure S3. TMEM150A and TMEM150B exhibit similar membrane diffusion dynamics.** (A) The mobile fraction (mean ± SD) of TMEM150A-GFP and TMEM150B-GFP, measured by FRAP. (B) The recovery times ($t_{1/2}$, mean ± SD) of TMEM150A-GFP and TMEM150B-GFP, measured by FRAP. n=17–19; ns, not significant.

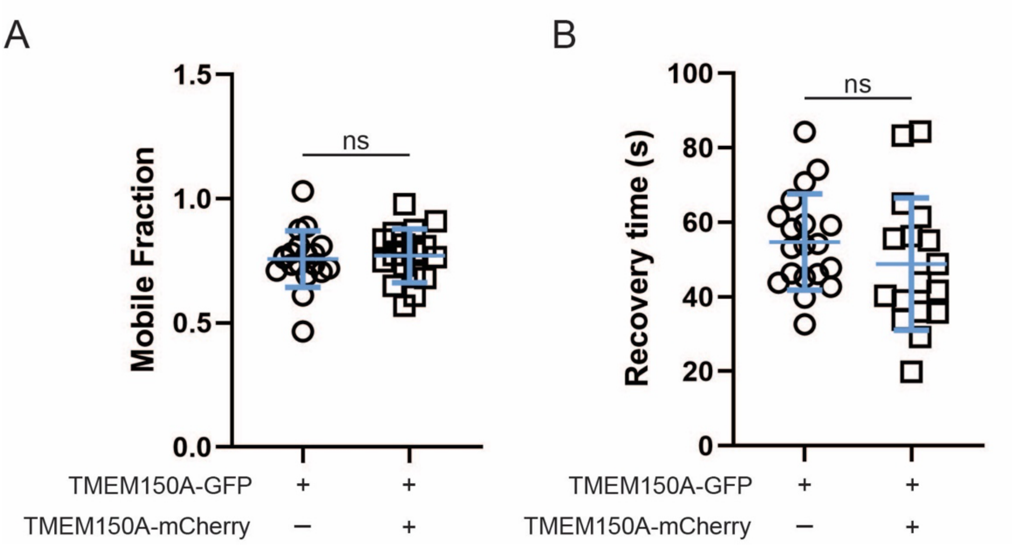

**Figure S4. FRAP reveals that the slow diffusion rate of TMEM150A is not due to oligomerization in the PM.** (A) Mobile fractions (mean ± SD) of TMEM150A-GFP when expressed alone or in the presence of TMEM150A-mCherry, measured by FRAP. (B) Recovery times ($t_{1/2}$, mean ± SD) of TMEM150A-GFP when expressed alone or in presence of TMEM150A-mCherry, measured by FRAP. Note that overexpression of TMEM150A-mCherry does not influence the mobile fraction or diffusion of TMEM150A-GFP, suggesting that substantial oligomerization does not occur between the two proteins. n=14–17; ns, not significant.

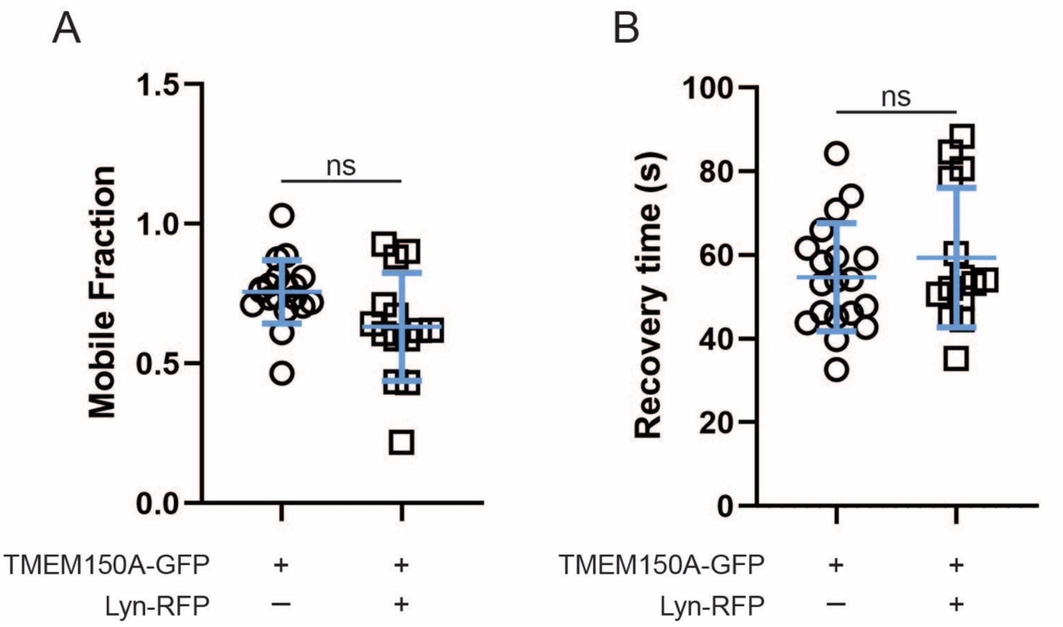

**Figure S5. TMEM150A-GFP does not engage in nonspecific interactions with Lyn, a different palmitoylated protein than EFR3B.** (A) Mobile fractions (mean ± SD) of TMEM150A-GFP when expressed alone or in presence of Lyn-RFP, a model palmitoylated and myristoylated membrane anchor. (B) Recovery times ($t_{1/2}$, mean ± SD) for TMEM150A-GFP in the presence of absence of Lyn-RFP. The mobile fraction and recovery time of TMEM150A are not influenced by the presence of Lyn-RFP, indicating that the diffusion properties of TMEM150A as measured by FRAP are not sensitive to overexpression of a palmitoylated protein unrelated to the PI4KIIIα complex. n=14–19; ns, not significant.

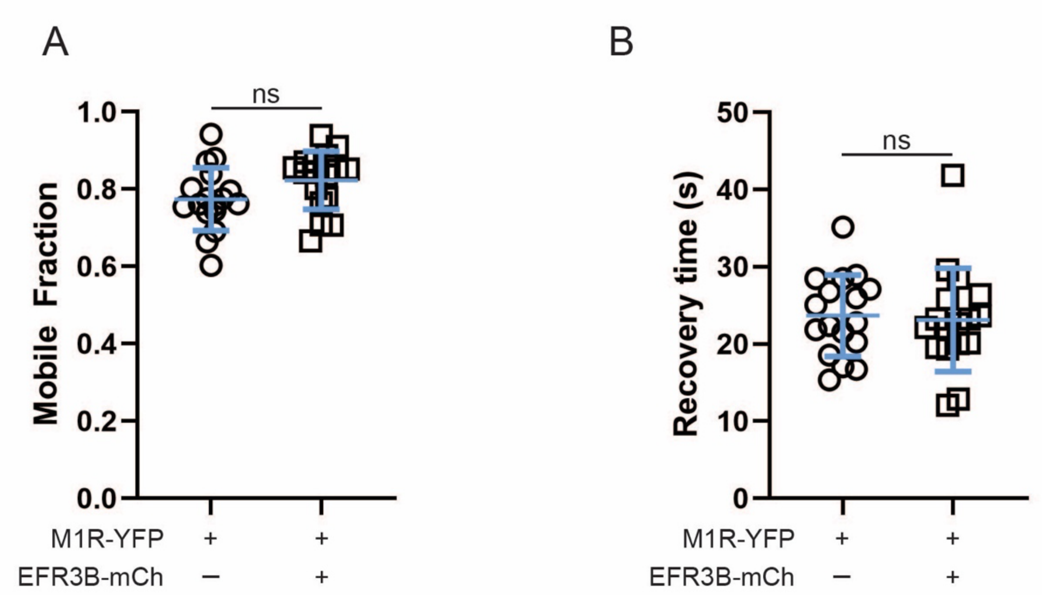

**Figure S6. EFR3B overexpression does not cause an increase in the diffusion of the muscarinic M1 receptor (M1R), a different transmembrane protein than TMEM150A.** (A) The mobile fraction (mean ± SD) of M1R-YFP in the presence of absence of EFR3B-mCherry measured by FRAP. (B) The recovery time ($t_{1/2}$, mean ± SD) of M1R-YFP in the presence and absence of EFR3B-mCherry measured by FRAP. Note that the mobile fraction and recovery time of M1R-YFP is not modified by the overexpression of EFR3B, suggesting that the effect of EFR3B on TMEM150A diffusion is specific. n=17; ns, not significant.

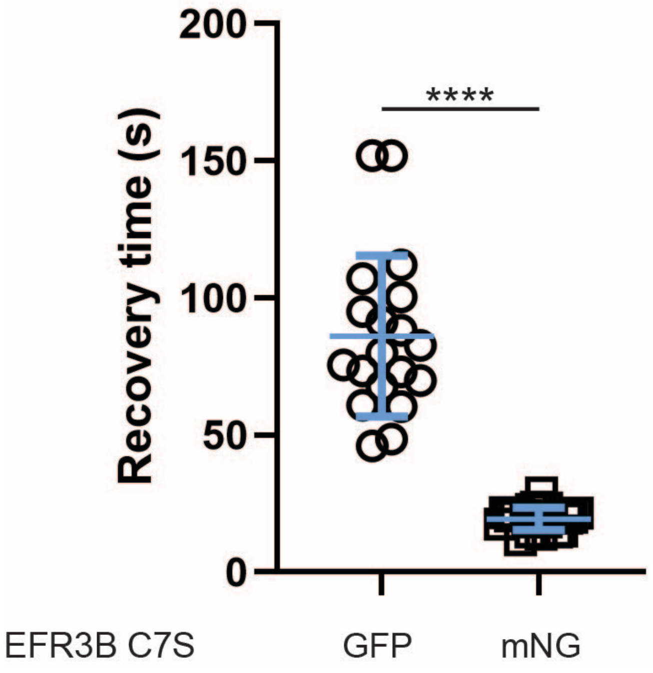

**Figure S7. Clustering of the EFR3B(C7S) mutant clustering modifies its membrane diffusion properties.** Recovery times (mean ± SD) of EFR3B(C7S)-GFP and EFR3B(C7S)-mNG, measured by FRAP. Note that the GFP-tagged construct, which forms clusters visible by confocal microscopy, exhibits much slower diffusion. By contrast, the mNG construct, which does not exhibit such clustering, results in diffusion properties similar to the other GFP-tagged CxS mutants (see Fig. 2E). n=19–20; ****, $p < 0.001$

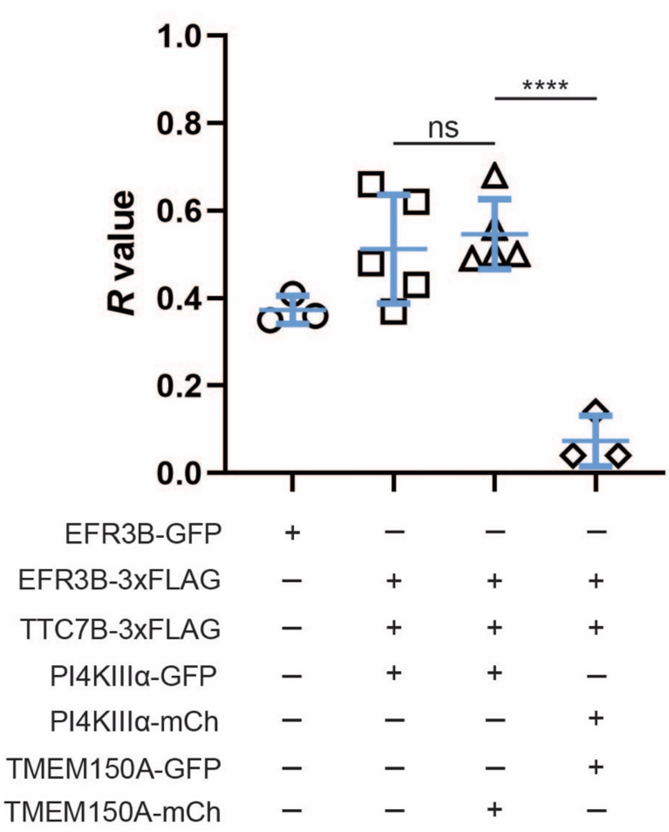

**Figure S8. The TTC7B-containing Complex I and TMEM150A-containing Complex II co-exist in the PM.** (A) *R* values (mean ± SD) from iDRM assays showing the fraction of retained GFP fluorescence from RBL-2H3 cells expressing the indicated combination of constructs, after a mild wash with 0.04% TX-100. Lane 1: EFR3B-GFP is moderately detergent resistant when expressed alone. Lanes 2–3: PI4KIIIα is highly detergent resistant in both the TTC7B-containing Complex I (lane 2) and TMEM150A-containing Complex II (lane 3). Lane 4: TMEM150A is highly sensitive to detergent even when Complex II is formed. n=3–5; ****, p<0.001; ns, not significant.

**Table S1. Sources of reagents, primers, antibodies, and plasmids.**

| Reagents | Source | Catalog number |
|---|---|---|
| DMEM | Corning | 10-017-CV |
| FBS | VWR | 45000-734 |
| Dialyzed FBS | VWR | 97065-302 |
| Penicillin/Streptomycin | VWR | 45000-652 |
| Gentamicin | Thermo Fisher | 15750078 |
| Transfectagro | VWR | 71002-816 |
| Lipofectamine 2000 | Invitrogen | 11668019 |
| cOmplete protease inhibitor | Millipore Sigma | 5056489001 |
| Fugene | Promega | E2312 |
| Phorbol 12,13-dibutyrate (PDB) | Sigma-Aldrich | P1269 |
| BCA assay | Pierce | 23225 |
| TCEP hydrochloride | Cayman Chemical Company | 14329 |
| N-ethylmaleimide | Alfa Aesar | 40526 |
| Hydroxylamine | Allied Chemical | 1789 |
| Methoxypolyethylene glycol maleimide (5 kDa) | Sigma-Aldrich | 63187 |
| Lactacystin | Santa Cruz Biotechnology | sc-3575 |
| Cycloheximide | Amresco | 94271 |
| Ezview Red Anti-FLAG M2 Affinity beads | Sigma-Aldrich | F2426 |
| 17-Octadecynoic acid (alk-16) | Cayman Chemical Company | 90270 |
| Cy5.5 azide | Click Chemistry Tools | 1059-1 |
| THPTA (tris-hydroxypropyltriazolylmethylamine) | Click Chemistry Tools | 1010 |
| Sodium L-ascorbate | Chem Impex | 01436 |
| Cupric sulfate | Mallinckrodt | 4844 |
| Clarity ECL reagent | Bio-Rad | 1705061 |
| Oxotremorine M | Santa Cruz | Sc-203656 |
| Atropine | TCI | A0550 |
| **Primers** | **Source** | **Sequence** |
| EFR3B C5S-3xFLAG | IDT | 5'-agggcaccgcagcagccagacacaccgtac-3'<br>5'-gtacggtgtgtctggctgctgcggtgccct-3' |
| EFR3B C7S-3xFLAG | IDT | 5'-ggtgtgtgtggcagctgcggtgccc-3'<br>5'-gggcaccgcagctgccacacacacc-3' |
| EFR3B C8S-3xFLAG | IDT | 5'-gtgtgtggctgcagcggtgcccttc-3'<br>5'-gaagggcaccgctgcagccacacac-3' |
| EFR3B C5,7S-3xFLAG | IDT | 5'-tgtgtctggctcctgcggtgcccttc-3'<br>5'-gaagggcaccgcaggagccagacaca-3' |
| EFR3B C5,8S-3xFLAG | IDT | 5'-tgtgtctggctcctgcggtgcccttc-3'<br>5'-agggcaccggagcagccagacacac-3' |

| | | |
|---|---|---|
| EFR3B C7,8S-3xFLAG | IDT | 5'-ggtgtgtgtggcagcagcggtgcccttc-3'<br>5'-gaagggcaccgctgctgccacacacacc-3' |
| EFR3B C5S-GFP/mCherry | Eurofins Genomics<br>IDT | 5’-catgtacggtgtgagtggctgctgcgg-3’<br>5’-agaggtaccgagtatacacacagatcaggaaacttcatctcatagac-3’ |
| EFR3B C7S-GFP/mCherry | IDT | 5’-cagagaattcaccatgtacggtgtgtgtggctcctgc-3’<br>5’-agaggtaccgagtatacacacagatcaggaaacttcatctcatagac-3’ |
| EFR3B C8S-GFP/mCherry | IDT | 5’-cagagaattcaccatgtacggtgtgtgtggctgctccg-3’<br>5’-agaggtaccgagtatacacacagatcaggaaacttcatctcatagac-3’ |
| EFR3B C5,7,8S-GFP/mCherry | Eurofins Genomics | 5’-cgaagggcaccgctgctgccactcacaccgtacatg-3’<br>5’-catgtacggtgtgagtggcagcagcggtgcccttcg-3’ |
| M1R-3xFLAG | Eurofins Genomics | 5’-aagcggccgcagccaccatgaacacctcagtgccccctgc-3’<br>5’-actagaatccgcattggcgggagggggtgc-3’ |
| **Antibodies** | **Source** | **Catalog number** |
| Rabbit anti-FLAG polyclonal | Sigma-Aldrich | F7425 |
| Mouse anti-FLAG M2 monoclonal | Millipore Sigma | F1804 |
| Anti-GFP | Takara | 632375 |
| Anti-GAPDH | Genetex | GTX78213 |
| Anti-Calnexin | Abcam | Ab22595 |
| Anti-rabbit-HRP | Bio-Rad | 1706515 |
| Anti-mouse-HRP | Bio-Rad | 1706516 |
| **Plasmids** | **Source** | |
| EFR3B-3xFLAG | Gift from the De Camilli lab | |
| EFR3B C5S-3xFLAG | This paper | |
| EFR3B C7S-3xFLAG | This paper | |
| EFR3B C8S-3xFLAG | This paper | |
| EFR3B C5,7S-3xFLAG | This paper | |
| EFR3B C5,8S-3xFLAG | This paper | |
| EFR3B C7,8S-3xFLAG | This paper | |
| EFR3B C5S-GFP | This paper | |
| EFR3B C7S-GFP | This paper | |
| EFR3B C7S-mNeonGreen | This paper | |
| EFR3B C8S-GFP | This paper | |
| EFR3B C5,7S-GFP | This paper | |
| EFR3B C5,8S-GFP | This paper | |
| EFR3B C7,8S-GFP | This paper | |

| | |
|---|---|
| EFR3B C5,7,8S-tdTomato | Gift from the De Camilli lab |
| EFR3B C5,7,8S-GFP | This paper |
| EFR3B C5,7,8S-3xFLAG | This paper |
| TMEM150A-GFP | Gift from the De Camilli lab |
| TMEM150A-mCherry | This paper |
| TTC7B-mCh | Gift from the De Camilli lab |
| GFP-PI4KIIIα | Gift from the De Camilli lab |
| iRFP-PH(PLCδ) | Gift from the De Camilli lab |
| M1R-YFP | Gift from the De Camilli lab |
| M1R-3xFLAG | This paper |
